# The beneficial rhizobacterium *Bacillus velezensis* acquires iron from roots via a type VII secretion system for colonization

**DOI:** 10.1101/2021.08.01.454677

**Authors:** Yunpeng Liu, Xia Shu, Lin Chen, Huihui Zhang, Haichao Feng, Xiting Sun, Qin Xiong, Guangqi Li, Weibing Xun, Zhihui Xu, Nan Zhang, Corné M.J. Pieterse, Qirong Shen, Ruifu Zhang

**Affiliations:** Key Laboratory of Agricultural Microbial Resources Collection and Preservation, Ministry of Agriculture and Rural Affairs, Institute of Agricultural Resources and Regional Planning, Chinese Academy of Agricultural Sciences, Beijing 100081, P. R. China; Jiangsu Provincial Key Lab for Organic Solid Waste Utilization, National Engineering Research Center for Organic-based Fertilizers, Jiangsu Collaborative Innovation Center for Solid Organic Waste Resource Utilization, Nanjing Agricultural University, Nanjing, 210095, P.R. China; Experimental Center of Forestry in North China, Chinese Academy of Forestry, Beijing 102300, P. R. China; Plant-Microbe Interactions, Department of Biology, Utrecht University, Utrecht, the Netherlands

**Author notes:** Corresponding author Yunpeng Liu, Ruifu Zhang. These authors contributed equally to this study (Yunpeng Liu, Xia Shu, Lin Chen). **Author contributions** Conceptualization, Y.L. and R.Z.; Formal Analysis, Y.L., L.C., X.Shu and G.L; Investigation, Y.L., X.Shu, H.Z. and X.Sun; Verification, H.F. and Q.X.; Writing-Original Draft, Y.L.; Writing-Review & Editing, Y.L., L.C., C.M.J.P. and R.Z.; Visualization, Y.L. and X.Shu; Supervision, W.X., Z.X., C.M.J.P. and R.Z.; Project Administration, N.Z. and Q.S.; Funding Acquisition, Y.L. and R.Z.

**Keywords:** Beneficial rhizobacteria, Type VII secretion system, iron, Bacillus, root colonization

## Abstract

Niche colonization is the key for bacterial adaptation to the environment, and competition for iron largely determines root colonization by rhizosphere microbes. Pathogenic and beneficial symbiotic bacteria use various unique secretion systems to support plant colonization or acquire limited resources from the environment. However, ubiquitous nonsymbiotic beneficial rhizobacteria have never been reported to use a unique secretion system to facilitate colonization. Here, we show that the type VII secretion system (T7SS) of the beneficial rhizobacterium *Bacillus velezensis* SQR9 contributes to root colonization. Knocking out T7SS and the major secreted protein YukE in SQR9 caused a significant decrease in root colonization. Moreover, the T7SS and YukE caused iron loss in plant roots in the early stage after inoculation, which contributed to root colonization by SQR9. Interestingly, purified YukE, but not inactivated YukE, could change the permeability of root cells. We speculated that secreted YukE might be directly inserted into the root cell membrane to cause iron leakage, indicating that the bacterial protein and root cell membrane interact directly. Moreover, a bacterial siderophore and the T7SS may be coordinately involved in iron acquisition by *B. velezensis* SQR9 for efficient root colonization. We showed that the beneficial rhizobacterium *B. velezensis* SQR9 could acquire iron from roots via the T7SS for rapid colonization. These findings provide the first insight into the function of the unique secretion system in nonsymbiotic beneficial rhizobacteria and reveal a novel mutualism in which plants and bacteria might share iron in a sequential manner.

## Introduction

Host-microbe interactions have drawn much research attention due to their importance in agricultural production and human health. Efficient colonization is a prerequisite for microbes to exert potential virulent or beneficial functions on their host (Arnaouteli et al., 2021). Soil microbial diversity is high, the competition among the microbes is strong, and rapid colonization is important for the occupation of nutrient-enriched niches on roots by soil bacteria (De Coninck et al., 2015; Haas and Défago, 2005; Lugtenberg et al., 2001). A group of important rhizoplane-colonizing bacteria, termed plant growth-promoting rhizobacteria (PGPRs), has been widely used in agricultural production due to the multifunctional plant-beneficial properties of its members (Bulgarelli et al., 2013; Siddiqui, 2006). These nonsymbiotic beneficial bacteria natively inhabit the rhizosphere and have been isolated, enriched and reapplied in the rhizosphere to benefit plants in a variety of ways, such as by suppressing soil-borne disease, regulating plant development, enhancing stress tolerance and increasing nutrient availability (Philippot et al., 2013).

Iron plays a critical role in plant growth, development and defense and is also a limited resource in the rhizosphere, competed for by microbes to support colonization and simultaneously fight against competitors (Harbort et al., 2020). Phytopathogens and plants always fight for iron to enhance their leading role in interactions (Herlihy et al., 2020), while beneficial rhizobacteria can enhance plant iron uptake. Paradoxically, sufficient iron is also required by beneficial rhizobacteria for colonization and for formation of biofilms on plant roots (Xu et al., 2019). In some cases, high iron levels may be toxic to plants due to the overproduction of reactive oxygen species (ROS) (Sun et al., 2021). Therefore, restriction of iron in plant roots, especially during rhizobacterium recognition, is reasonable but has never been reported.

For competition and colonization, a range of animal- or plant-associated pathogenic and symbiotic bacteria have developed unique secretion systems to transfer certain proteins from the cytoplasm across the membrane or a hydrophobic structure to the extracellular space, thereby interacting with the hosts (Chang et al., 2014; Costa et al., 2015). However, nonsymbiotic beneficial rhizobacteria have never been known to use a unique secretion system to support their colonization. Among these systems, the type VII secretion system (T7SS) is the only unique secretion system in gram-positive bacteria (Abdallah, 2007). It was discovered for its role in tuberculosis caused by *Mycobacterium bovis*. Recent studies revealed that EsxA, the core protein secreted by the T7SS of *Mycobacterium tuberculosis*, plays an essential role in phagosome rupture and bacterial cytosolic translocation within host macrophages by insertion into the phagosome membrane (Conrad et al., 2017; De Leon et al., 2012; Zhang et al., 2020). T7SS is also widely conserved in *Bacillus* species (Abdallah, 2007), the most widely used PGPRs in agricultural production (Arnaouteli et al., 2021). *Bacillus* species possess the *yuk/yue* operon encoding the T7SS for secretion of YukE, which is homologous to EsxA in *Mycobacterium* (Baptista et al., 2013; Huppert et al., 2014; Sysoeva et al., 2014). It has been reported that the T7SS operon in *Bacillus subtilis* is adequate for secretion under the regulation of a global regulator (Baptista et al., 2013; Huppert et al., 2014; Sysoeva et al., 2014). However, the biological and bioecological functions of the *Bacillus* T7SS are unknown. In particular, is this T7SS in beneficial rhizobacteria involved in the interaction between bacteria and the plant host? And if so, what roles does it play in that process?

*Bacillus velezensis* SQR9 is a well-studied rhizobacterium that colonizes plant roots and benefits plants by promoting growth and controlling disease (Li et al., 2021; Liu et al., 2017; Wu et al., 2018; Zhang et al., 2021). It uses siderophores and specific iron ABC transporters to acquire iron, thereby supporting biofilm formation and root colonization (Xu et al., 2019). In this study, we found that the T7SS and its secreted protein YukE caused iron loss from plant roots, and this effect then contributed to root colonization by *B. velezensis* SQR9. We further found that the T7SS was induced by iron, thus theoretically supporting the model that T7SS function is amplified in the root-iron reduction process and plays an important role in rapid root colonization by *Bacillus* strains that are beneficial to plants. Our study is the first to report a unique secretion system that contributes to root colonization by a nonsymbiotic beneficial rhizobacterium, which may represent a novel mechanism in plant-microbe interactions in which bacterial molecules directly change the permeability of root cells.

## Results

### A type VII secretion system contributes to root colonization by the rhizobacterium *B. velezensis* SQR9

Genome analysis of *B. velezensis* SQR9 (China General Microbiology Culture Collection Center, CGMCC accession no. 5808) revealed that the *yuk* and *yue* clusters, *yukE*, *yukD*, *yukC*, *yukAB*, *yueB*, *yueC*, *yueD*, and *yueE* (V529_31490) constitute the T7SS (Fig. 1A). Based on the collected RNA-seq results of 72 SQR9 samples from 5 independent projects, we found high transcriptional correlations of *yueD*, *yueC*, *yueB*, *yukAB* and *yukC* (Fig. 1B). Transcription analysis indicated that *yukE*, *yukD*, *yukC*, *yukAB*, *yueB*, *yueC*, and *yueD* are transcribed as polycistronic mRNAs (Fig. S1), suggesting that they may all contribute to the function of the T7SS. YukE was reported to be the only protein secreted by the T7SS in *Bacillus* (Baptista et al., 2013; Huppert et al., 2014; Sysoeva et al., 2014). Our analysis demonstrates that YukE is one of the most highly transcribed genes (top 5 among 3873 genes, Table S1), indicating its important role in *B. velezensis*.

**Figure 1.**
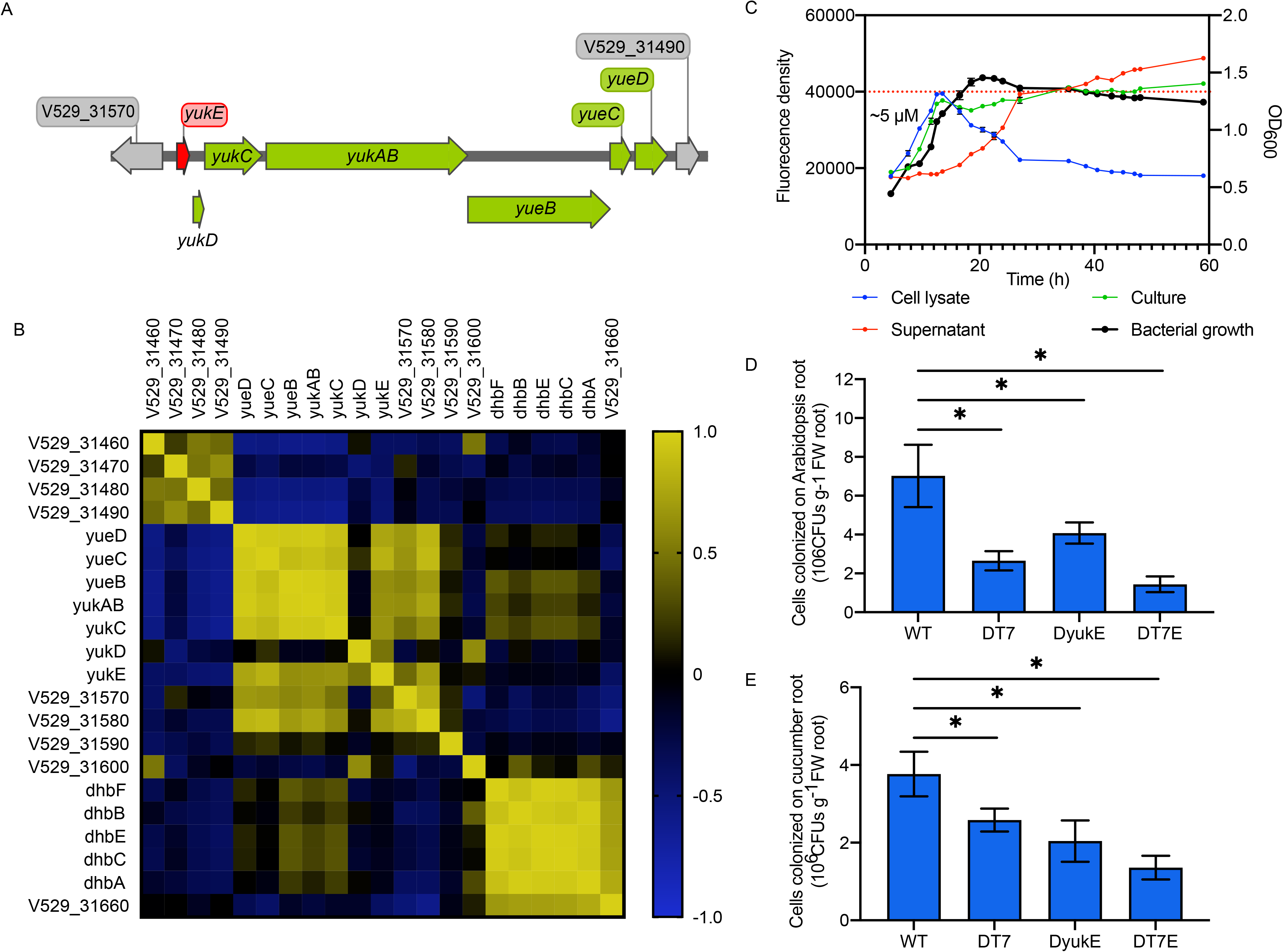
Contribution of the *Bacillus velezensis* SQR9 type VII secretion system (T7SS) to root colonization. (A) Schematic illustration of the *yuk/yue* operon encoding the T7SS. The red arrow indicates the gene encoding the secreted WXG100 family protein YukE, and the green arrow indicates the genes encoding the export machinery. (B) Transcriptional correlation of the *yuk/yue* operon based on 72 RNA-seq results. Pearson correlation was calculated; colors indicate the R values of the transcriptional correlations between each pair of genes: yellow color indicates high correlation, and blue color indicates low correlation. (C) Content of YukE in the supernatant and cell lysate. *B. velezensis* SQR9 YukE was fused with GFP at the N-terminus in situ, and for detection, the excitation and emission wavelengths were set at 488 nm and 533 nm, respectively. The whole culture, supernatant and cell lysate from one sample were measured, and the O_600_ was recorded to indicate the cell density. Measurement of each time point included six independent replicates. Error bars indicate the standard errors. The dotted line indicates the fluorescence density of 5 μM purified YukE-GFP. (D and E) Colonization of roots of *Arabidopsis* and cucumber grown in a hydroponic environment. Colonization was performed within 1/2 MS medium, and the colonized bacterial cells were diluted and spread on plates for counting. Fifteen-day-old Arabidopsis Col-0 or “Chinese long 9930” cucumber with two fully expanded true leaves was used for the colonization assay. Error bars indicate the standard errors. Six replicates were included for each strain. Asterisks indicate significant differences based on Student’s t test (P<0.05). The experiment was repeated three times with similar results.

To investigate the effect of the T7SS on root-rhizobacterium interactions, the T7SS secreted protein-coding gene *yukE*, the T7SS component-coding region (*yukD-yueD*) and the whole T7SS (*yukD-yueD* and *yukE*) were deleted, and the mutants, designated DyukE, DT7 and DT7E, respectively, were obtained. A GFP tag was fused to the N-terminus of YukE in situ, which had no effect on the secretion property of YukE at a concentration of 5 μM in medium (Fig. 1C). Root colonization analysis of *Arabidopsis* and cucumber showed that at 3 days post inoculation, deletion of *yukE* or the T7SS significantly reduced root colonization by *B. velezensis* SQR9 on both *Arabidopsis* and cucumber (Fig. 1D and 1E). Deletion of the T7SS and *yukE* together (DT7E) also significantly reduced the root colonization efficiency (Fig. 1D and 1E). Comparison of these mutants with wild-type (WT) SQR9 in terms of growth, biofilm formation and motility, which are the factors affecting root colonization, showed no significant differences (Fig. S2, Fig. S3, Fig. S4). These results suggest that the T7SS and YukE contribute to root colonization by interacting with plants rather than by affecting bacterial properties.

### *B. velezensis* SQR9 decreased the root iron content in the early colonization stage via the T7SS and YukE

To explore the mechanism by which the T7SS and YukE affect root colonization by *B. velezensis* SQR9, RNA-seq was performed with *Arabidopsis* roots responding to WT *B. velezensis* SQR9, DT7, DyukE, DT7E, purified YukE and inactivated YukE protein at 1 h, 3 h, 6 h and 24 h post inoculation or treatment. These bacteria were inoculated into the rhizosphere at a final concentration of 10^6^ cells/mL, and YukE (Fig. S5) or inactivated YukE was applied to the rhizosphere at a final concentration of 5 μM. Differentially expressed genes (DEGs) in YukE-treated and inactivated YukE-treated roots indicated that YukE enriched Gene Ontology (GO) terms associated with iron binding and heme binding; moreover, functional annotation by DAVID (DAVID Bioinformatics Resources 6.8) also enriched the keywords iron, heme and oxidoreductase (Fig. 2A). Furthermore, GO analysis showed that the unique DEGs in the DT7, DyukE or DT7E treatment in comparison with the WT SQR9 treatment were all significantly enriched in the defense response (Fig. S6). Since the plant defense response occurs downstream of iron homeostasis signaling, we hypothesized that the T7SS and YukE may affect the plant iron content. We then performed Q-PCR to determine whether the iron acquisition genes were affected in roots. The key component for acquiring iron in Arabidopsis roots is shown in Fig. S7A. At 6 h and 24 h, the expression of FIT and MYB72, regulators of iron acquisition gene expression (Herlihy et al., 2020), was differentially affected by WT SQR9 and the T7SS-related mutant strains (Fig. S7B and C). Expression of the direct regulator FIT in the DT7, DyukE and DT7E treatments was significantly higher than that in the WT SQR9 treatment (Fig. S7B and C). Consistent with this, the downstream genes *IRT1* and *FRO2* showed the same expression profiles (Fig. S7D and E). Expression of *FER1*, encoding an iron storage protein, was also significantly induced by WT *B. velezensis* SQR9, while the expression of this gene in the DT7, DyukE and DT7E treatments was significantly lower than that in WT SQR9 treatment (Fig. S7F). These results showed that the plant gene expression profile after WT SQR9 treatment was the opposite of the iron deficiency response, while mutation of the T7SS compromised this effect. These results suggest that the SQR9 T7SS contributed to changes in plant iron homeostasis.

**Figure 2.**
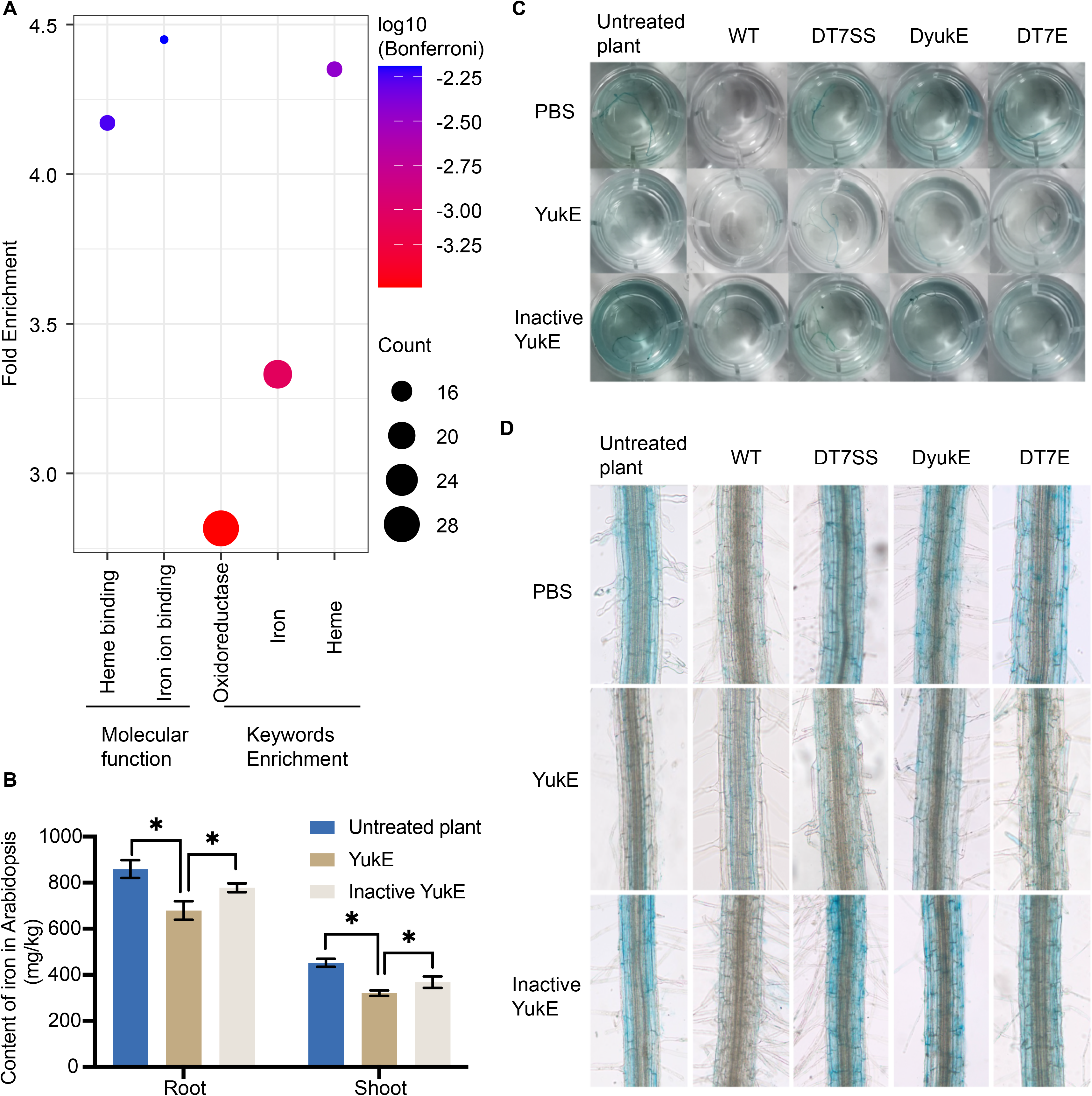
Effect of the *Bacillus velezensis* SQR9 type VII secretion system (T7SS) on the transcription of plant iron acquisition genes. (A) Functional enrichment of the DEGs in the YukE treatment but not in the inactivated-YukE treatment. Briefly, 5 μM purified YukE or inactivated YukE was added to the plant medium, and RNA-seq was performed at 1 h, 3 h, 6 h and 24 h post treatment. The fragments per kilobase of exon per million fragments mapped (FPKM) value of each gene in the YukE or inactivated-YukE treatment was compared with that of untreated plants to identify the upregulated and downregulated DEGs. Then, DEGs in the YukE and inactivated-YukE treatments were compared to identify the DEGs present in the YukE treatment but not in the inactivated-YukE treatment. These DEGs were then subjected to functional enrichment analysis using DAVID (https://david.ncifcrf.gov) (Huang et al., 2009). (B) ICP-MS measurement of the iron content in roots and shoots of Arabidopsis treated with YukE or inactivated YukE. Asterisks indicate significant differences based on Student’s t test (P<0.05). (C) Image of Perls staining of Arabidopsis inoculated with *B. velezensis* SQR9 or treated with YukE. YukE or inactivated YukE was added to the rhizosphere at a final concentration of 5 μM; *B. velezensis* SQR9 or the derived strains were inoculated at a final concentration of 10^6^ cells/mL. The roots were cut after 24 h for Perls staining. (D) Microscopic view of Perls staining of the root iron content.

To further confirm this finding, we measured the plant root iron content by inductively coupled plasma mass spectrometry (ICP-MS), which showed that YukE reduced the root and shoot iron content, while inactivated YukE did not have this effect (Fig. 2B). Perls staining assays at 24 h, 48 h and 72 h post inoculation of the bacteria showed that WT *B. velezensis* SQR9 dramatically decreased the root iron content at 24 h post inoculation (Fig. 2C, 2D and Fig. S8). Interestingly, deletion of T7SS, *yukE* or both compromised the SQR9-induced root iron loss (Fig. 2C, 2D and Fig. S8), and exogenous purified YukE rescued this effect (Fig. 2C 2D). These results indicated that *B. velezensis* SQR9 could significantly decrease the plant iron content via the T7SS and YukE. However, at 48 h and 72 h post inoculation, neither WT SQR9 nor these mutants affected the root iron content (Fig. S8), suggesting that the decrease in root iron content caused by SQR9 T7SS occurred only in the early stage of root colonization.

### Excess rhizosphere iron or YukE rescues the colonization defect of the SQR9 T7SS mutants

Iron is a critical rhizosphere nutrient that is competed for among rhizosphere microbes and by microbes and plants. As deletion of T7SS or the secreted protein YukE in *B. velezensis* SQR9 resulted in root colonization deficiency, while WT SQR9 caused root iron loss, we hypothesize that *B. velezensis* SQR9 may use the T7SS to secrete YukE to acquire iron from root cells in the early interaction stage to facilitate colonization; in this case, supplementation with exogenous YukE or iron should rescue the colonization defect of DT7, DyukE and DT7E. To verify this hypothesis, a root colonization assay was performed with modified Murashige and Skoog (MS) medium containing 2-fold iron (180 μM). As speculated, in the iron-rich MS medium, DT7, DyukE and DT7E did not show differences in root colonization, while in the regular MS medium, DT7SS, DyukE and DT7E all showed significantly lower root colonization (Fig. 3A). This result indicated that excess iron rescued the root colonization defect of these mutants (Fig. 3A). In parallel, colonization by WT SQR9 and the mutants on *Arabidopsis* roots in iron-free medium was tested. Both WT SQR9 and the T7SS mutants showed dramatically reduced root colonization in iron-free medium, but the iron limitation amplified the colonization defect of the T7SS mutants compared with that of WT SQR9 (Fig. 3A). Moreover, with exogenous YukE protein (5 μM), the root colonization defect of DT7SS, DyukE and DT7E was completely rescued (Fig. 3A). These results supported the hypothesis that YukE functions in the acquisition of iron from plant roots to benefit *B. velezensis* SQR9 colonization.

**Figure 3.**
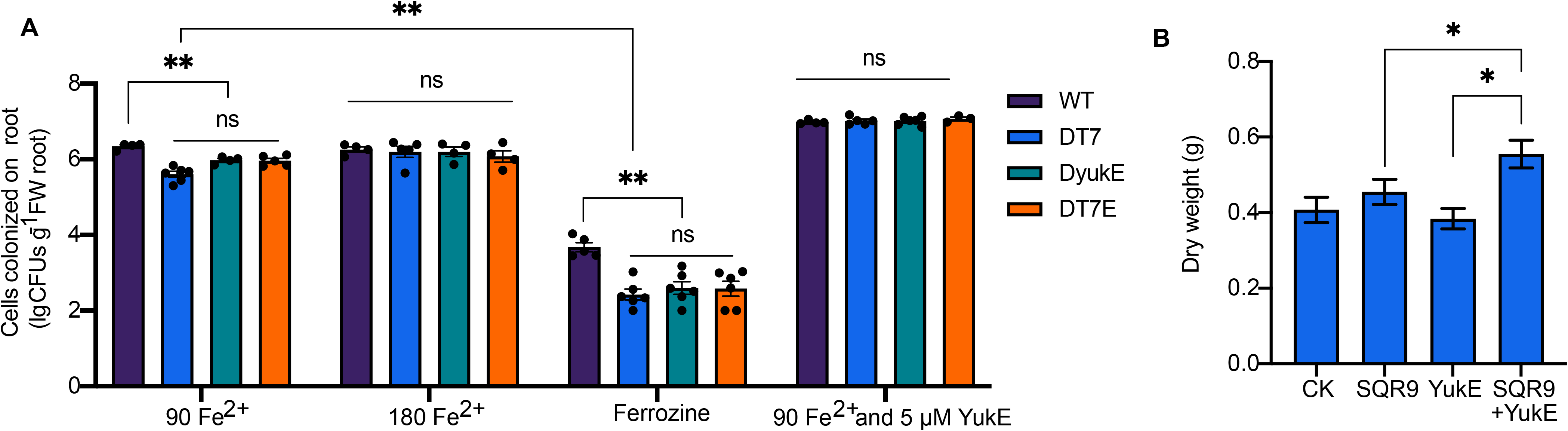
YukE functions in root colonization by reducing the plant iron content and enhances the growth promotion effect of *B. velezensis* SQR9. (A) Colonization by *B. velezensis* SQR9 on *Arabidopsis* roots under hydroponic conditions. *Arabidopsis* was grown in iron-free MS medium and was manually supplied with FeCl2. Ferrozine at a final concentration of 300 μM was supplied to generate iron-free conditions. YukE was supplied at a final concentration of 5 μM. Six independent replicates were included for each treatment. The experiment was performed twice and showed similar results. Error bars indicate standard errors. (B) Growth promotion of cucumber by *B. velezensis* SQR9 in pots. Cucumber was inoculated by dipping into a cell suspension for 1 day before transplantation into soil. YukE was supplied in the cell suspension at a final concentration of 5 μM. Ten independent replicates were included for each treatment.

To exclude the possibility that the rescue phenotype may be accidentally caused by changes in the growth or biofilm formation of the SQR9 T7SS mutants under low- or high-iron conditions, we then evaluated the growth and biofilm formation of DT7SS, DyukE, DT7E and WT SQR9 in medium with different iron concentrations. The results showed that although excess iron (100 μM) in MSgg medium enhanced biofilm formation of all these strains, low iron reduced biofilm formation (Fig. S9), and no differences were observed between different strains at these tested iron concentrations (Fig. S9). Differences in the iron concentration did not influence the growth of these strains in either MS or MSgg medium (Fig. S10), and excess YukE protein had no effect on the growth and biofilm formation of these strains (Fig. S11 and S12). These results indicate that a higher iron level could enhance biofilm formation and thus root colonization by *B. velezensis* SQR9, and the decrease in the root iron content was essential for the function of the T7SS and YukE in root colonization.

Since the plant roots showed a decrease in iron content post YukE treatment, we wondered whether YukE is disadvantageous to plant growth. Interestingly, we found that excess YukE significantly enhanced the growth promotion effect of SQR9 (Fig. 3B). This result indicated that the effect of YukE on the decrease in root iron content was temporary and did not negatively affect plant growth, which was also explained by the fact that the decrease in iron content was no longer observed after 24 h (Fig. S8).

### YukE may be directly inserted into the root cell membrane to cause iron leakage

To explore the mechanisms of the YukE-induced decrease in the root iron content, *B. velezensis* SQR9 and the YukE protein were labeled with RFP and GFP, respectively, and the strain SQR9-RFP/YukE-GFP was obtained to observe the location of YukE on plant roots. Interestingly, it was observed that most of the bacterial cells in the rhizosphere completely secreted YukE at 24 h post inoculation (Fig. 4A), and the secreted YukE was enriched and located on the root cell surface before bacterial colonization (Fig. 4A). This result suggested that YukE might function on the root surface.

**Figure 4.**
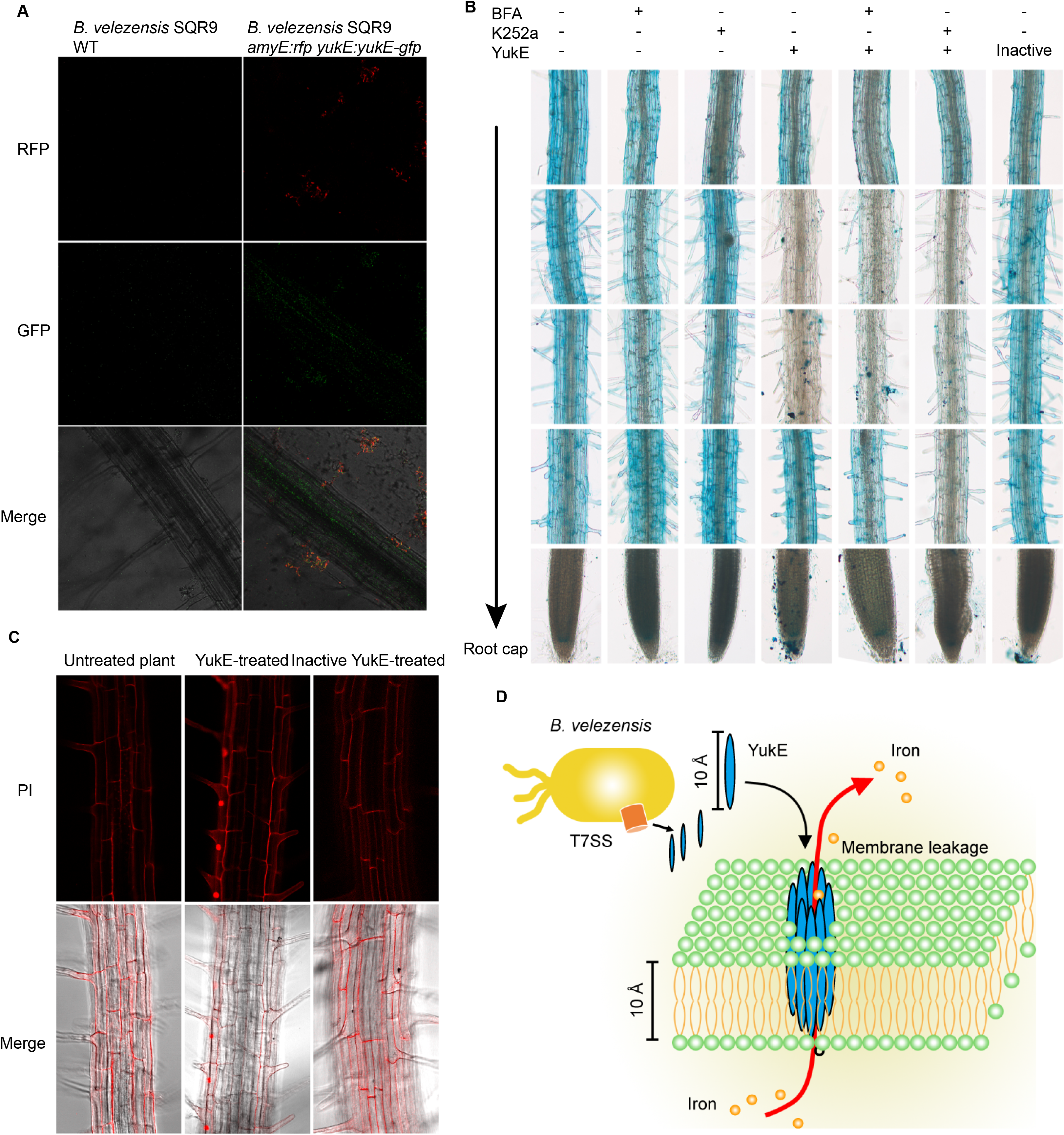
YukE function and location on the root surface. (A) Location of YukE in the rhizosphere. YukE of *B. velezensis* SQR9 was fused with GFP, and bacterial cells were labeled with RFP under the control of the constitutive promoter P43. Fluorescence was observed by confocal laser scanning microscopy (CLSM). The images were taken at 24 h post inoculation. (B) Effect of K252a and BFA on the iron-decreasing capability of YukE. YukE was supplied at a final concentration of 5 μM, and K252a and BFA were supplied at final concentrations of 1 μM and 100 μg/mL, respectively. YukE inactivated in a boiling water bath was included as a negative control. Three replicates showed similar results. (C) Effect of YukE on the staining of root cell nuclei. Roots were treated with YukE or mock control for 24 h and then stained with 10 μg/mL PI in the dark and visualized by CLSM. Three replicates showed similar results. (D) Model for the role of YukE in the decrease in root iron content. The predicted structure of YukE consists of two helices with a helix-turn-helix motif (WXG motif). The protein existed as a dimer in vivo and in vitro. The length of the protein is 10.1 Å. The thickness of the phospholipid bilayer is approximately 10 Å. YukE may be inserted into the phospholipid bilayer and cause iron leakage from root cells.

YukE may act as a signal that is sensed by root receptors to inhibit plant iron acquisition. To confirm or exclude this possibility, K252a, a general kinase inhibitor blocking the potential YukE-sensing process of plants, and brefeldin A (BFA), a general inhibitor of exocytosis in plant cells, were applied individually to determine whether they blocked the decrease in plant iron content caused by YukE. Neither K252a nor BFA alone affected the plant iron content (Fig. 4B), and the iron-decreasing effect of YukE was not compromised in the presence of K252a or BFA, indicating that a plant root cell surface kinase sensor or exocytosis was not involved in YukE-induced root iron reduction (Fig. 4B). These results suggest that the decrease in the root iron content caused by YukE might be signal transduction independent, implying that YukE may decrease the root iron content in a direct manner.

The *M. tuberculosis* T7SS cargo protein EsxA, which is homologous and structurally similar to YukE, has been reported to be inserted into the host membrane and form a pore, causing leakage of the phagolysosome (Ma et al., 2015; Zhang et al., 2020). We wondered whether YukE may reduce the plant root iron content directly by insertion into the root cell membrane to cause iron leakage. To confirm whether YukE changed the root cell membrane structure, YukE-treated plant roots were stained with the general dye propidium iodide (PI), and the results showed that the cell nuclei of YukE-treated roots, but not untreated roots, could be stained by PI (Fig. 4C). PI can stain plant cell nuclei only when the plant cell membrane permeability is changed. In addition, we did not observe plant root cell death by microscopy (data not shown). In conclusion, we propose that YukE may interact with the root cell membrane and cause leakage of iron in the early colonization stage, thus facilitating rapid root colonization (Fig. 4D).

Structure analysis of YukE showed no membrane spanning region. But the inside of the YukE structure built based on the model of EsxA in *M. tuberculosis* showed enriched hydrophobic residues between the 4 helixes (Fig. S14A, B and C). We proposed YukE might have a change structure when present in membrane that expose the hydrophobic side out. Moreover, based on structure comparison, we found the crystal form of *Geobacillus thermodenitrificans* EsxA (3ZBH), which is homologues of YukE (56.7% identity), was in an asymmetric unit including four homo dimers (Fig. S14D, E and F). The asymmetric unit formed a pore-like structure in side-by-side structure (Fig. S14D, E and F). This structure proposed that YukE homodimers could form an advanced structure bound with their hydrophilic side to each other thus expose the hydrophobic side out. The T7SS and YukE are conserved in *Bacillus*, including both plant-associated and animal-associated species. Sequence analysis of YukE of different *Bacillus* species indicated that the YukE proteins were divided into two groups (Fig. S13). Interestingly, one group, which included *B. velezensis* and *B. subtilis*, was associated with plant roots, while the other group, which included *B. anthracis* and *B. thuringiensis*, was associated with animals. The animal-associated group also showed a close phylogenetic relationship with *Staphylococcus aureus,* a common animal pathogenic bacterium (Fig. S13). These results support the idea that YukE might be an important factor deciding the host of *Bacillus* species (Fig. S13).

### The switch of T7SS- and siderophore-dependent iron acquisition in SQR9 is under iron regulation

In addition to the T7SS in *Bacillus*, siderophore-dependent iron acquisition is also an important strategy via which bacteria acquire environmental iron. *B. velezensis* SQR9 produces bacillibactin (BB, synthesized by DhbA, DhbC, DhdE, DhdB and DhbF; Fig. 1B) as a siderophore to bind iron in the environment, and the BB-bound iron is then transported into bacterial cells by FeuABC (Xu et al., 2019). To distinguish T7SS- and siderophore-dependent iron acquisition, SQR9 mutants with BB or FeuA deficiency were obtained, but they did not compromise the ability to decrease the root iron content (Fig. 5A). In contrast to the BB-based strategy, isothermal titration calorimetry (ITC) and native gel Perls staining showed that iron-free YukE could not bind iron (Fig. 5B and C). These results indicated that YukE reduced the plant root iron content but could not bind iron, while the BB-Feu system did not reduce the plant root iron content. Therefore, the T7SS- and siderophore-dependent iron acquisition systems should have a clear division of labor—while siderophores function in a low-iron environment to compete for iron with other microbes, the T7SS represents an additional strategy by which rhizosphere *Bacillus* species temporally acquire a large amount of iron to occupy niches as fast as possible. We then evaluated how the T7SS- and siderophore-based strategies were affected by the environmental iron level. Indeed, the results showed that T7SS expression under high levels of iron (5 μM and 50 μM) was significantly higher than that under low levels of iron (0 Fe) (Fig. 5D), while in siderophore strategy, the transcriptional regulator Btr was induced under low levels of iron (Fig. 5E). The intracellular or extracellular level of YukE was consistent with the RT-PCR results (Fig. 5FGHI), indicating that YukE mainly functions under regular iron conditions but not for competition in low-iron conditions. The finding also suggested that YukE secretion by *B. velezensis* and the decrease in root iron content were in a positive feedback loop. When root iron leakage occurred, secretion and production of YukE were induced and caused more iron leakage (Fig. 5J).

**Figure 5.**
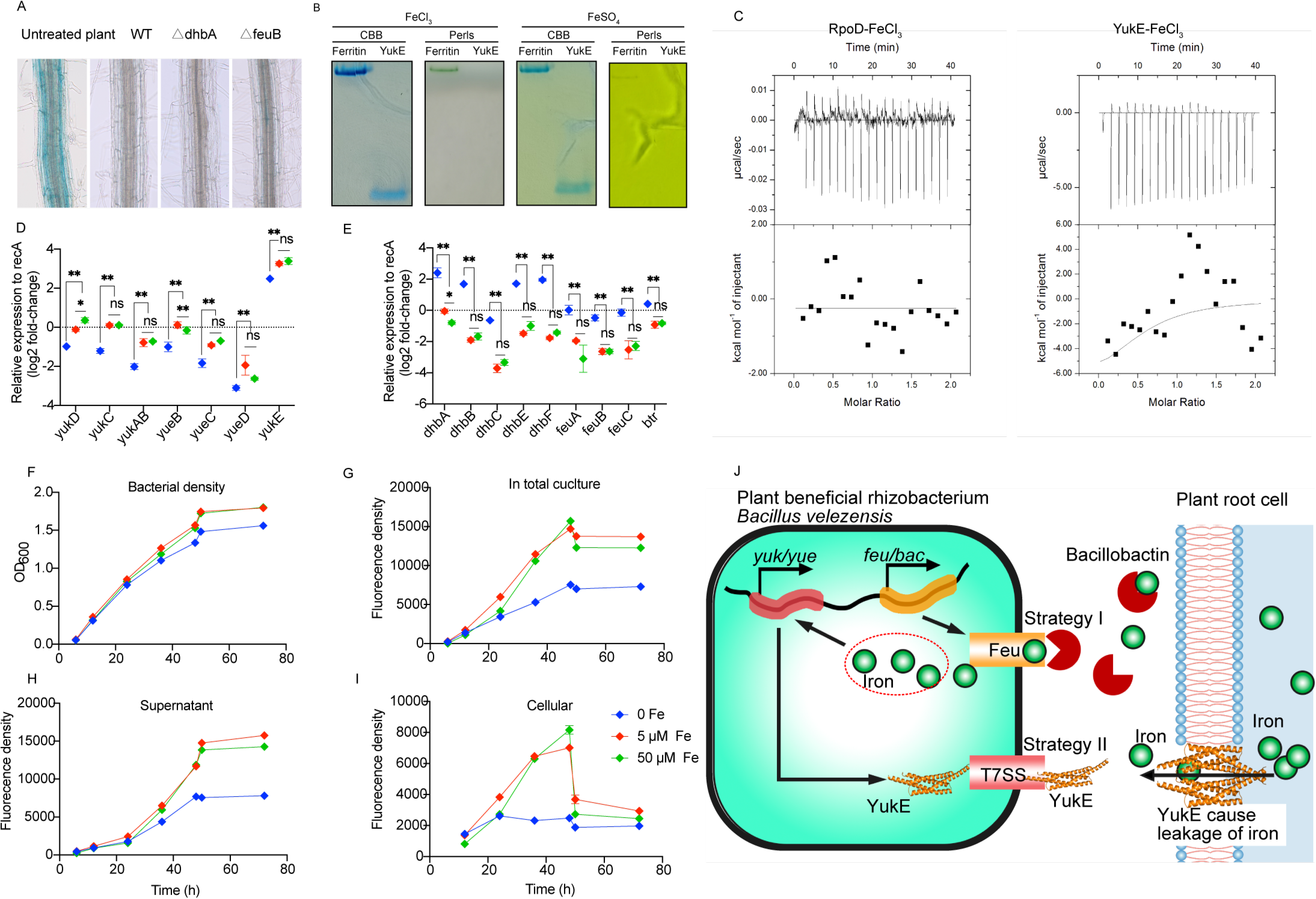
Comparison of the iron acquisition strategy with the siderophore-Feu system in *B. velezensis* SQR9. (A) Microscopic view of Perls staining of the root iron content. *Arabidopsis* was inoculated with *B. velezensis* or the siderophore- or Feu-deficient mutant. (B) Native binding of Fe^3+^ and Fe^2+^ to YukE. The gel with His-tag-free YukE was incubated with Fe^3+^ or Fe^2+^ and then stained with Coomassie brilliant blue (CCB) or Perls’ blue. Ferritin was included as a positive control that could bind iron. (C) ITC for detection of the binding between YukE and Fe^3+^. His-tag-free YukE was used to exclude the binding of the 6His tag and Fe^3+^. (D-E) qRT-PCR for measuring the expression of genes in the T7SS (B) and siderophore-dependent iron acquisition strategy (C). (F-I) Effect of iron conditions on YukE secretion by *B. velezensis* SQR9. YukE was fused with GFP in vivo. F, Bacterial density recorded with OD600. G, YukE production recorded as the fluorescence density of the total culture. H, YukE secretion from cells recorded as the fluorescence density of the supernatant of the culture. I, YukE in cells recorded as the fluorescence density of the lysate of bacterial cells. Error bars indicate the standard errors. (J) Model of the two iron acquisition strategies in *B. velezensis* SQR9.

## Discussion

Unique bacterial secretion systems are crucial for adaption to the environment. Since the T7SS was discovered in *Bacillus*, the most important plant-beneficial microbe group, its biological and ecological significance has never been investigated. In the present study, we found that the *Bacillus* T7SS and the secreted protein YukE facilitated rapid root colonization by the beneficial rhizobacterium *B. velezensis* SQR9 via acquisition of iron from plant roots in the early colonization stage. We further found that YukE caused a change in root cell membrane permeability. Based on our results and the reported function in *Mycobacterium*, we further proposed that YukE might be inserted into the root cell membrane and cause iron leakage from root cells. We further illustrated the possible coordinating functions between the T7SS and siderophores in iron acquisition. These findings provide the first insight into T7SS function in a beneficial bacterium and reveal a novel mutualism in which plants and bacteria might share iron sequentially, thus benefiting both sides.

Gram-negative bacteria have developed a variety of unique secretion systems because their bilayer cell membrane structure limits protein transmembrane transport by a single membrane-spanning system (such as SEC and TAT) (Chang et al., 2014). However, the unique secretion system in gram-positive bacteria includes only the T7SS and is poorly understood. The T7SS has been found in only Actinobacteria and Firmicutes thus far and is also known as an ESX system because of its function in exporting the small proteins ESAT-6 and CFP-10 (EsxA and EsxB) in *Mycobacterium* (Simeone et al., 2009). Five T7SS clusters, termed ESX-1, ESX-2, ESX-3, ESX-4 and ESX-5, have been discovered in *Mycobacterium*, but only one is present in *Bacillus* (Abdallah, 2007). The *Bacillus* T7SS cluster is homologous to ESX-4, which is the simplest ESX (Abdallah, 2007). Evolutionary analysis has shown that the T7SS cluster in *Bacillus* might be the ancestor of all T7SSs discovered in bacteria (Newton-Foot et al., 2016). Evolution of the T7SS from certain monoderm bacteria such as *Bacillus* to that from species harboring a mycomembrane-like outer membrane containing mycolic acids, such as *Mycobacterium* species, also coincided with the evolution of prokaryotes from monoderm gram-positive bacteria to bilayer membrane-harboring gram-negative bacteria (Abdallah, 2007). This evidence suggests that the T7SS in *Bacillus* might be the most ancient unique secretion system in the bacterial kingdom. However, its biological function has never been understood before this study. Therefore, this study may have significance for the discovery of the drivers of the evolution of T7SS and even other unique secretion systems. It also helps elucidate the evolutionary divergence of ancient bacteria, such as the *B. subtilis* group and *B. anthracis* group.

In *Bacillus*, YukE is homologous to EsxA, a WXG100 family protein in the tuberculosis-causing pathogen *M. tuberculosis*. EsxA could be inserted into the lysosome membrane and contribute to lysosome lysis by *M. tuberculosis,* thereby helping with invasion from lysosomes into the cytoplasm of macrophages (Conrad et al., 2017; Ma et al., 2015; Zhang et al., 2020). We proposed that YukE contributes to iron competition by causing root cell iron leakage but not as a root-sensed molecule that affects plant iron acquisition. This hypothesis was based on three results: 1) YukE’s root iron-decreasing effect could not be abolished by the general kinase inhibitor K252a; 2) supplying YukE facilitated PI staining of the cell nucleus, while staining of the nucleus was not observed in the absence of YukE; 3) the WT strain SQR9 had more advantages in root colonization than the *yukE* or T7SS mutant in iron-free medium, in which condition the root is the only iron resource for the bacterium. This kind of direct interaction with the root cell membrane has never been proposed for rhizobacteria previously. In interaction between macrophage and *Legionella pneumophila*, Isaac et al. has found similar model that a type IV secretion system (T4SS) secreted protein, MavN, could insert into host membranes to mediate iron acquisition (Isaac et al., 2015). Different with that MavN secreted by T4SS from *L. pneumophila* might directly binding to iron (Isaac et al., 2015), in vitro experiment present in this study showed that YukE secreted by T7SS from *B. velezensis* could not bind iron directly (Fig 5B and 5C). Therefore, we supposed T7SS might coordinate with the siderophore strategy to complete the iron-acquisition process. Nonetheless, further study is needed to show whether YukE causes cell leakage and the structure that YukE forms in the membrane. In addition, this study is not the only one showing the relation between the T7SS and iron. Casabona et al. found that deletion of an essential component of the T7SS from *S. aureus* RN6390 caused an iron starvation response in the bacterium; moreover, its T7SS was transcriptionally regulated by iron (Casabona et al., 2017).

Plant pathogens use diverse strategies to steal iron from plants (Herlihy et al., 2020), but beneficial bacteria have never been found to decrease the iron content in plants previously. The T7SS is very highly conserved in these beneficial *Bacillus* strains; in other words, the iron-decreasing effect in the early colonization stage should generally occur during this interaction. It would be interesting to understand how plants adapt to the decrease in iron content and why they did not evolve to resist the effect. As shown above and in a previous study, an ROS burst takes place along with colonization by *B. velezensis* in the very early stage (Zhang et al., 2021), and the ROS can react with cytoplasmic iron to produce hydroxyl radical (·OH), which is a kind of ROS that is deleterious to cells (Ravet et al., 2009), through the Fenton reaction. The runaway Fenton reaction leads to accumulation of ·OH, which damages cells and could even cause ferroptosis in some cases (Apel and Hirt, 2004; Herlihy et al., 2020). Excess iron and ROS are considered to be major factors in plant immunity-triggered cell death (ferroptosis) (Dangol et al., 2019; Herlihy et al., 2020). It has been reported that a chemical that triggers iron and ROS accumulation in plants leads to plant cell death when plants are challenged with non-cell death-triggering bacteria under regular iron conditions (Dangol et al., 2019). Due to the Fenton reaction, we speculated free iron in the cytoplasm is harmful for plants in the very early stage of root colonization by *B. velezensis* (Dangol et al., 2019; Reyt et al., 2015). In contrast, iron is an important factor for biofilm formation of *B. velezensis* (Xu et al., 2019). In this study, we found that the timings of iron reduction and ROS production were consistent (Fig. S8) (Zhang et al., 2021). We suggest that decreasing iron in the early colonization stage may benefit plants by contributing to the avoidance of unexpected ferroptosis. When the ROS burst ended, the iron content-reducing effect was terminated by an unknown mechanism before 24 h post inoculation. Moreover, bacteria also regulate plant immunity by binding iron; for instance, siderophores from bacteria regulate plant immunity by binding iron (Dellagi et al., 2009). This may also partially explain the contribution of the iron-decreasing effect of the T7SS to colonization.

The iron-decreasing effect was observed in only the first 24 h (Fig. S8). At 48 h post inoculation, all treatments with *B. velezensis* SQR9 or the mutants had no effect on root iron content in comparison with that in untreated plants (Fig. S8). Moreover, YukE alone did not show a negative effect on plant growth (Fig. 3B); moreover, it enhanced the growth-promoting effect of *B. velezensis* SQR9 (Fig. 3B). In conclusion, the YukE-induced decrease in root iron content within a limited duration is a win-win interaction, in contrast to the unilateral effect of pathogenic bacteria (Khan et al., 2018). However, how the decrease in iron content was terminated was not answered in this study. We found that the secretion of YukE from *B. velezensis* cells occurred soon after inoculation (YukE was completely secreted from cells in 24 h) and caused iron leakage from root cells (Fig. 4A). As the iron leakage from root cells increased, the secretion and transcription of YukE were induced (Fig. 5D and 5H), thereby causing further iron leakage via feedback. This result indicated that termination of the decrease in root iron content might not have been due to the termination of secretion or production. As we suggested that YukE might be inserted into the root cell membrane and cause leakage, we believe that termination might be achieved by the degradation of YukE by the bacterium or plant or by the disassociation of YukE from the root cell membrane. Further study on the interaction between YukE and the phospholipid bilayer will answer this question.

Overall, this is the first report showing that a unique secretion system contributes to colonization by beneficial bacteria. It is also the first report showing that plants share iron with rhizobacteria for mutual beneficial interactions.

### Prospects

Further investigation of the interaction of YukE and the lipid membrane may reveal the interaction mechanisms of rhizobacterial proteins and the plant cell membrane. Does YukE form a pore in the root cell membrane specific for iron leakage? What is the structural basis for its function on the root cell membrane? How is the leakage terminated after the first 24 h? Moreover, it is necessary to determine what other components leak from root cells under the effect of YukE. How does *Bacillus* remain the first beneficiary of the root iron leakage effect in a complex soil environment? Answering these questions will greatly expand the knowledge of the interaction between plant roots and rhizobacteria. It would also theoretically support research on tuberculosis and the evolution of bacterial secretion systems.

