## Supplemental materials for "The beneficial rhizobacterium *Bacillus velezensis* acquires iron from roots via a type VII secretion system for colonization"

[*]Corresponding author

### SUPPLEMENTARY MATERIALS

### EXPERIMENTAL MATERIALS

#### Arabidopsis

*Arabidopsis thaliana* ecotype Col-0 was used in this study. Arabidopsis seeds were surface sterilized with 2% (vol/vol) NaClO. The sterile seeds were placed in petri dishes containing 1/2 Murashige and Skoog (MS) medium with 2% (wt/vol) sucrose and 0.8% (wt/vol) agar and vernalized for 2 days at 4°C in darkness. Then, seeds on petri dishes were cultured under 16-h light and 8-h dark cycles at 22°C for one week, and the seedlings were then transferred to new petri dishes containing 1/2 MS medium with 0.5% (wt/vol) sucrose and 1.5% (wt/vol) agar and cultured for one more week for experimental use.

**Cucumber**

The 'Chinese long' cucumber inbred line 9930 was used in this study. The cucumber seeds were surface sterilized with 75% (vol/vol) ethanol and then with 2% (vol/vol) NaClO. The seeds were planted into axenic tissue culture bottles containing vermiculite and allowed to germinate and grow for 4 days in a growth chamber at 28°C with a 16-h light and 8-h dark photoperiod. The seedlings were then transplanted into 50-mL flasks containing 35 mL of sterile liquid (1/2 sucrose-free MS medium) to submerge the seedling roots in the medium, with one seedling in each flask. The seedlings were cultured in these flasks for 3 weeks. The MS medium was replaced every other day during the growth period. The hydroponic system was gently shaken (50 rpm) for 2 h each day on a shaker.

***Bacillus velezensis* SQR9**

*Bacillus velezensis* SQR9 (China General Microbiology Culture Collection Center, CGMCC accession number 5808) and the derived strains were grown at 30°C on Luria-Bertani (LB) medium (10 g/L peptone, 5 g/L yeast extract and 5 g/L NaCl) agar plates. MSgg medium (5 mM potassium phosphate (pH 7), 100 mM 3-(N-morpholino)propanesulfonic acid (pH 7), 2 μM $MgCl_2$, 700 μM $CaCl_2$, 50 μM $MnCl_2$, 50 μM $FeCl_3$, 1 μM $ZnCl_2$, 2 μM thiamine, 0.5% glycerol, 0.5% glutamate, 50 μg/mL tryptophan and 50 μg/mL phenylalanine) was used for biofilm formation (Liu et al., 2020). When necessary, antibiotics were added at the following final concentrations: chloramphenicol (Cm) at 5 mg/L, zeocin at 20 mg/L, spectinomycin (Spc) at 100 mg/L, and kanamycin (Kan) at 30 mg/L.

### METHOD DETAILS

**Root colonization assay**

For Arabidopsis, twelve-day-old *A. thaliana* Col-0 plants were transferred from 1/2 MS agar plates to 6-well plates containing liquid 1/2 MS with 0.5% sucrose (3 seedlings per well). After 7 days, the medium was replaced with differentially modified 1/2 MS medium without sucrose and $FeSO_4$ for the experiment. Generally, the MS medium without sucrose and Fe used in the colonization period was modified to include 300 μM ferrozine, 90 μM $FeSO_4$ or 180 μM $FeSO_4$ to generate an iron concentration gradient. Ferrozine has high affinity for iron and was used to capture iron, thereby producing iron-free conditions. For the experiment with YukE, the purified YukE protein was added to the rhizosphere at a final concentration of 5 μM. Moreover, wild-type SQR9 and the mutated strains were inoculated into each well at a final optical density at 600 nm ($OD_{600}$) of 0.01. The root samples were harvested 2 days after treatment and washed three times with water. The samples were placed in 2-mL tubes containing 3 zirconium beads, weighed and suspended in 1 mL of 0.1 M PBS using a Mixer Mill MM400 (Retsch, Haan, Germany) with a frequency of 30 beats per second. Bacterial densities were assessed by plating dilutions of the samples onto LB agar medium. The plates were incubated at 30°C for 12 h, and then, the numbers of colony forming units (CFUs) were determined. Each treatment included six replicates from independent wells.

**Biofilm formation assay**

The biofilm formation assay was performed as described by Liu et al. (Liu et al., 2020). *B. velezensis* SQR9 and its derived strains were cultured in LB medium until the $OD_{600}$ reached 1.0. The bacterial cells were collected by centrifugation at 8000 rpm. The pellets were washed with sterile water three times and suspended in MSgg medium at an $OD_{600}$ of 1.0. Biofilms were formed in 48-well microtiter plates. Each well was filled with 1 mL of MSgg medium inoculated with a 10-μL suspension of *B. velezensis* SQR9 or the derived strains. The negative control contained only the corresponding culture medium. For biofilm formation under unique conditions, such as in iron-free, low-iron, and YukE-containing media, the final concentration used was the same as that in the colonization experiment. Every biofilm formation condition included 4 replicates.

**Motility assay**

The swarming assay was performed following the method described by Inoue et al. (Inoue et al., 2007). Briefly, *B. velezensis* SQR9 and its derived strains were cultured in LB medium until the $OD_{600}$ reached 0.8. The cultures were inoculated on petri dishes containing semisolid LB medium (0.5% (w/v) glucose and 0.6% (w/v) Eiken agar (Eiken, Nogi-machi, Japan)) by sterilized toothpicks. The petri dishes were incubated at 37°C for 4-6 h to allow the bacteria to swim. Then, the water content of the medium in the petri dishes was reduced with dry air to terminate the swimming process. Subsequently, the petri dish was incubated overnight at room temperature. The swarming circles on the petri dishes were observed and recorded after

24 h.

**Protein purification**

Exogenous expression and purification of 6His-YukE, 6His-YukE-GFP, and 6His-GFP were performed in an *E. coli* BL21(DE3)/pET29a system. The DNA fragments encoding the full-length YukE or GFP were amplified from SQR9 genomic DNA or pNW33N-gfp, respectively. The PCR products were cloned into the expression plasmid pET29a (+) (Novagen, Madison, U.S.A.) by a one-step cloning kit (Vazyme Biotech Co., Ltd.). The reconstructed plasmid was verified by DNA sequencing. The plasmid was introduced into *E. coli* BL21(DE3) for expression. The expressed proteins contained an N-terminal 6His tag. The obtained strains were grown in LB medium at 37°C with shaking at 200 rpm. When the $OD_{600}$ reached 0.5, the growth temperature was lowered to 16°C. After another 30-min incubation, protein expression was induced by adding 0.03 mM isopropyl-b-D-thiogalactopyranoside. The incubation was continued at 16°C with shaking at 200 rpm overnight prior to cell harvest by centrifugation at 8,000 × g for 10 min. The cell pellets were resuspended and disrupted by sonication (2-s pulse on, 3-s pulse off). The cellular lysates were centrifuged at 20,000 × g for 50 min. Finally, for further clarification, the lysates were passed through a 0.22-μm filter (Millipore), followed by purification with His-affinity resin chromatography. The purified proteins were collected and stored in PBS at -80°C.

The protein used for ITC of metal ions and protein was prepared following the method described by Si et al. (Si et al., 2017). For the His-tag-free YukE preparation, the expressed protein contained an N-terminal 6His tag and a TEV protease recognition site. The preliminarily purified proteins were collected and treated with TEV protease at 4°C overnight to remove the 6His tag. The 6His-tag-free YukE was purified by His-affinity resin chromatography for digestion to remove the proteins without a 6His-tag and TEV protease sire. Finally, the purified His-free proteins were collected and dialyzed overnight at 4°C in Buffer I, which contained 250 mM EDTA, 5 mM o-phenanthroline, 50 mM HEPES (pH 6.0), 150 mM NaCl and 10% (v/v) glycerol, followed by three dialysis steps in Buffer I without EDTA and o-phenanthroline.

**Genetic assay for *B. velezensis***

Gene deletion mutants were constructed using unmarked genetic manipulation based on multiple gene deletions by relying on a counterselectable marker, *pheS* (Feng et al., 2018). All these mutants were verified by PCR and sequencing.

We also labeled YukE with GFP at the N-terminus in vivo. Essential antibiotic resistance genes were included as selectable markers. Detailed information on the primers used for genetic manipulation is shown in Supplementary Table S3. The secretion of YukE was verified by western blotting using an anti-GFP antibody.

**Bacterial fluorescence density detection**

The fluorescence density was determined with the total culture, supernatant, and lysate of *B. velezensis* SQR9/YukE-GFP cells to determine the YukE concentration. The standard curve was obtained by detection of the purified 6His-YukE-GFP protein expressed with the pET29a-*E. coli* BL21 protein expression system. The fluorescence density was measured by an Infinite M200 PRO (Tecan, Männedorf, Switzerland) controlled by Tecan i-control software (v1.12). Six replicates were included.

**Bacterial growth measurement**

Growth curves of SQR9 and the mutant strains in LB medium or under the same conditions as the biofilm formation assay in modified MSgg medium were measured. Measurement of $OD_{600}$ was performed every hour using the Bioscreen C system. Six replicates were included for each treatment.

**Iron content measurement**

Roots of *Arabidopsis* grown in 6-well plates were inoculated with *B. velezensis* SQR9 or the derived mutant strain or treated with YukE. *B. velezensis* SQR9 was inoculated at a final concentration of $10^7$ cells/mL. YukE was added to the rhizosphere at a final concentration of 5 μM. At 24 h, 48 h and 72 h post inoculation or treatment, roots were collected for iron content measurement. For evaluation of the effect of YukE on the plant iron content, K252a and brefeldin A (BFA) dissolved in DMSO were added together with YukE if necessary at final concentrations of 1 μM and 100 μg/mL, respectively.

For Perls staining, roots were vacuum soaked in Perls staining solution containing 4% (w/v) K-ferrocyanide and 4% (v/v) HCl for 15 min and washed three times with deionized water to terminate the reaction and completely remove the solution (Hsiao et al., 2017; Lei et al., 2014). The formation of a blue pigment indicated the accumulation of iron. The results were observed and imaged using an Olympus light microscope under a 10× objective lens. The intensity of the staining was quantitatively analyzed by ImageJ. An equal square of the image was converted and evaluated for integrated density.

Iron content measurement was also performed by using inductively coupled plasma mass spectrometry (ICP-MS) as described previously (Xu et al., 2019). Briefly, plant tissues were harvested from 21-day-old plants and washed with deionized water three times to exclude Fe contamination from the medium. Subsequently, the tissues were dried at 70°C for 3 days and digested with $HNO_3$/$HClO_4$ (4:1, v/v). The resultant solution was subjected to ICP-MS. Every treatment included 3 replicates, and each replicate was pooled from 24 plants that were cultured in a 6-well plate.

**Western blot assay**

SQR9/*yukE-gfp* cells grown in LB medium for 48 h were harvested to collect the total cellular protein. After centrifugation and resuspension in PBS, 100 μM lysozymes were added to the cell suspension, and the mixture was incubated for 30 min to degrade the cell wall. Then, the cells were disrupted by sonication (2-s pulse on, 3-s pulse off). Subsequently, a TCA precipitation assay was performed to obtain the proteins. Proteins were separated by SDS-polyacrylamide gel electrophoresis (PAGE) and detected by Western blotting. Equal amounts of protein were loaded into each lane of an SDS–12.5% polyacrylamide gel. The protein was transferred to a cellulose nitrate membrane using an SPJ-1000Atransfer system (Pyxis). The membrane was sealed by 5% skim milk (v/v). The primary mouse GFP antibody used to detect YukE-GFP and standard anti-mouse antibody-horseradish peroxidase conjugate were from Sangon Biotech. Visualization was performed using BeyoECL Moon (Beyotime). The Western blots were scanned by using a ChemiDoc™ Imaging System (Bio-Rad).

**RNA extraction**

Root total RNA for qRT-PCR and RNA-Seq analyses was extracted using the RNeasy Plant Mini Kit (Qiagen, Hilden, Germany). Bacterial total RNA was extracted using the Bacterial RNA Kit (OMEGA, Biotek, USA). Plant total RNA was extracted using the RNeasy Plant Mini Kit (Qiagen, Hilden, Germany). The extracted RNA was evaluated on a 1% agarose gel, and the concentration and quality (A260/A280) were determined by a NanoDrop ND-2000 spectrophotometer (NanoDrop, Wilmington, DE, U.S.A.).

**Root RNA-seq**

Two-week-old *Arabidopsis* grown in six-well plates containing 3 mL of 1/2 MS medium was inoculated with *B. velezensis* SQR9 WT, DT7, DyukE or DT7E or supplied with purified YukE or heat-inactivated YukE. All inoculants or additives were dissolved in sterile water. Untreated plants were included as a control. At 1 h, 3 h, 6 h and 24 h post treatment, roots from the control and all the treated plants were cut and rapidly placed in liquid nitrogen for subsequent RNA extraction. Then, the extracted RNA was used for library preparation and sequencing.

The library was prepared and sequenced on an Illumina HiSeq 4000, and 150-bp paired-end reads were generated at the Beijing Allwegene Technology Company, Beijing, China. Clean reads were obtained after quality control and mapped to the reference genome using TopHat2. RNA-Seq data were normalized to the FPKM (fragments per kilobase of exon per million fragments mapped). Sequence data were deposited in the Sequence Read Archive (SRA) with the accession number PRJNA649312.

**Functional enrichment analysis**

The p value for the comparison of the gene expression was calculated using a negative binomial distribution-based test, the FDR (p adjust) was calculated using BH (Benjamini-Hochberg) correction and the differentially expressed genes (DEGs) were screened based on FDR < 0.05. The unique DEGs in the WT *B. velezensis* SQR9 treatment in comparison with each of the mutant treatments, the unique DEGs in each of the mutant treatments in comparison with the wild-type SQR9 treatment or the unique DEGs in YukE treatment in comparison with the inactivated-YukE treatment were recorded and subjected to DAVID (DAVID Bioinformatics Resources 6.8, NIAID/NIH) analysis for functional annotation. Gene Ontology (GO) enrichment, keyword enrichment, pathway enrichment and functional categories were all analyzed. The enrichment was evaluated with the p value and FDR by the Benjamini method. The enriched terms were screened with FDR<0.01.

**qRT-PCR**

The RNA samples were reverse transcribed to cDNA using the PrimeScript™ RT reagent Kit with gDNA Eraser (TaKaRa, Dalian, China). SYBR® Premix Ex Taq™ (Takara, Dalian, China) was used for qRT-PCR with a QuantStudio 6 Flex (Applied Biosystems, Foster City,

CA, USA). The following PCR program was used: cDNA was denatured for 30 s at 95°C, followed by 40 cycles of 5 s at 95°C and 34 s at 60°C. ACTIN2 was used as an internal reference (Ahmad et al. 2014). The reference genes used for bacteria and plants were *recA* and *actin2*, respectively. The specificity of the amplification was verified by melting curve analysis and agarose gel electrophoresis. Each treatment included four independent replicates.

**PI staining**

For propidium iodide (PI) staining, aseptic culture of 21-day-old *A. thaliana* Col-0 was treated with 5 μM YukE or inactivated YukE protein for 24 h and washed three times with deionized water. The roots were completely immersed in 10 μg/mL PI (purchased from Sigma-Aldrich) under dark conditions. The whole process was gentle to avoid damage to the roots. After the samples were allowed to stand for 10 min, the PI wavelength segments (488 nm excitation wavelength, 630 nm emission wavelength) were selected under a Zeiss laser confocal microscope for observation and photography.

**ITC**

The interaction between the protein and the metal ion was measured by isothermal titration calorimetry (ITC) following the method described by Wang et al. (Wang et al., 2015) and Si et al. (Si et al., 2017). ITC was performed using an ITC200 (Microcal) titration calorimeter at 25°C. The 6His-tag and metal ion were removed from the protein prior to determination. All ligands were prepared with the same solvent as the proteins.

For metal ion titration into the protein, 300 μL of 50 μM protein was placed in the sample cell and titrated with aliquots of ligand solution at 500 mM in the injector syringe. After a stable baseline had been achieved, 60 μL of metal ion solution ($FeCl_3$) was added with a total of 20 injections into the sample cell until the protein sample was saturated with metal ions.

**Staining of iron-binding proteins**

Ferritin (purchased from Sigma-Aldrich) and purified YukE protein were resolved by 12% native PAGE after incubation with 0.1 M $FeSO_4$ or $FeCl_3$ for 1 h at room temperature. The gel was then stained with $K_3[Fe(CN)_6]$ or $K_4Fe(CN)_6$ for 10 min in the dark and destained with 10% trichloroacetic acid/methanol solution. Then, the cells were stained with Coomassie blue after imaging. Horse spleen ferritin was used as a positive control (Saraswathi et al., 2009).

**Phylogenic analysis**

Phylogenic analysis of YukE in different *Bacillus* species was performed. Twenty-two YukE sequences from *Bacillus* and closely related species were collected, and the natural niches of the bacteria were recorded. The alignment was performed with ClustalW algorithms using MEGAX. The sequence logo was made by Weblogo (http://weblogo.berkeley.edu/logo.cgi). The phylogenetic analysis was inferred by using the maximum likelihood method based on the Tamura-Nei model. The initial tree for the heuristic search was obtained automatically by applying the neighbor-joining and BioNJ algorithms to a matrix of pairwise distances estimated using the maximum composite likelihood (MCL) approach and then selecting the topology with the superior log likelihood value. The analysis involved 22 nucleotide sequences. All positions containing gaps and missing data were eliminated. The phylogeny was tested by 500 bootstrap replications.

**CLSM**

The pNW33N plasmid carrying RFP was introduced into *B. velezensis* SQR9, and YukE was fused with GFP at the N-terminus. To view the position of *B. velezensis* SQR9 and the secreted YukE in the rhizosphere, 15-day-old Arabidopsis seedlings were transferred from a 6-well plate to a laser confocal dish and inoculated with the bacterium at a final concentration of $10^7$ cells/ml. The laser confocal dish with the inoculated Arabidopsis was placed under a CLSM system (Zeiss LSM880). The green and red fluorescence was monitored. The objective used was a Zeiss EC Plan-Apochromat 20 x/0.8 M27. The excitation wavelength and emission wavelength for GFP were 488 nm and 510 nm, respectively. The excitation wavelength and emission wavelength for RFP were 561 nm and 583 nm, respectively. More than twenty fields of vision were observed and showed similar results.

**Statistical analysis**

For intercorrelation analysis of genes in *B. velezensis* SQR9, RNA-seq data from 72 samples of *B. velezensis* SQR9 with FPKM values were collected. The FPKM values were then normalized by the sum and transformed to relative log expression (RLE) values. The Pearson correlation of the transcription between each gene pair was calculated by R (v3.6).

The data from different treatments were subjected to analysis of variance. Duncan's multiple range test ($P < 0.05$) was employed to determine differences among means and was performed using the R (v3.6) agricolae package. Pairwise statistical significance was analyzed by t tests; “*” and “**” indicate significant differences ($P<0.05$ and $P<0.01$, respectively). Analysis of variance was performed with Duncan’s multiple test; “*” and “**”, represent $P<0.05$, $P<0.01$, $P<0.001$ and $P<0.0001$, respectively. For analysis of the significance of the gene expression difference and GO enrichment, the P value was also adjusted by the Bonferroni method. Error bars in figures indicate the standard errors.

**SUPPLEMENTARY FIGURE CAPTIONS**

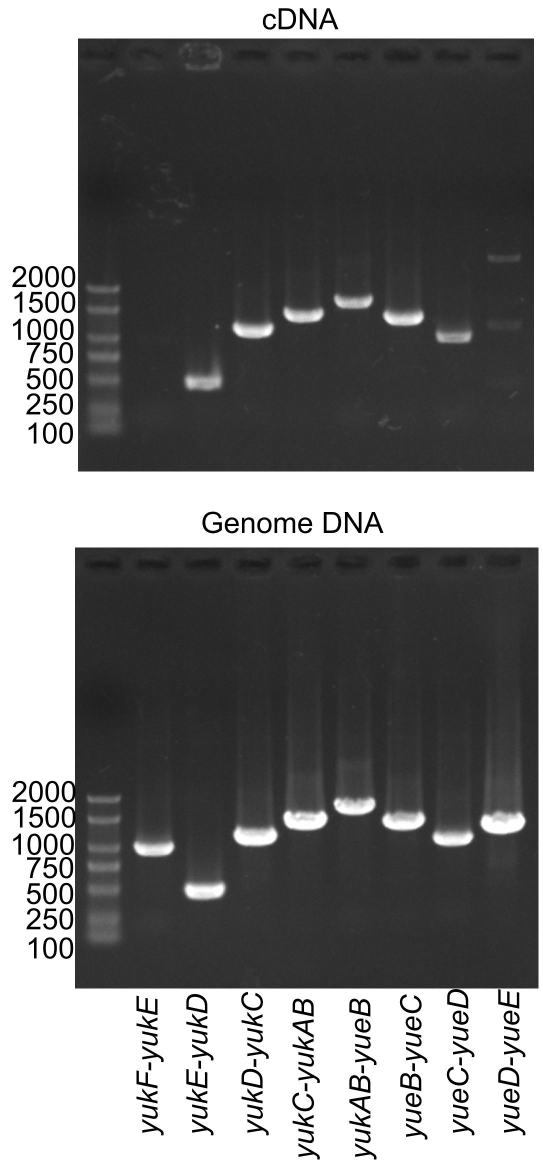


Fig. S1 Cross-gene PCR for operon verification. Cross-gene PCR was performed using cDNA and gDNA of *B. velezensis* SQR9. The *yuk* and *yue* clusters and the flanking region were located on the *B. velezensis* SQR9 genome in the order *yukF* (V529_31570), *yukE*, *yukD*, *yukC*, *yukAB*, *yueB*, *yueC*, *yueD*, *yueE*. *yukF-yukE* cross-gene PCR was included as a negative control. The estimated lengths of the PCR products from left to right were 1003 bp, 513 bp, 1147 bp, 1417 bp, 1716 bp, 1374 bp, 1070 bp and 1270 bp.

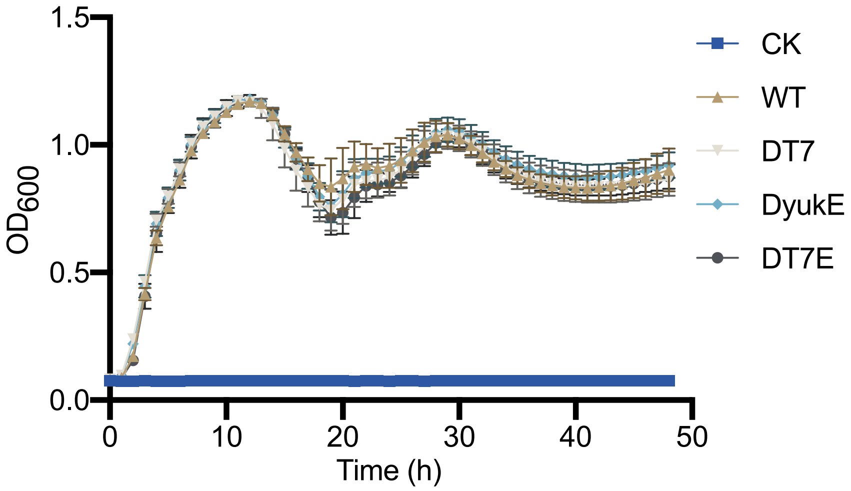


Fig. S2 Growth curve of *B. velezensis* SQR9 and the derived mutant strains in liquid LB medium. Growth of wild-type *B. velezensis* SQR9 and the mutants DT7, DyukE and DT7E in LB medium was measured by Bioscreen C. CK indicates medium without inoculation. Error bars indicate the standard error calculated from 6 independent replicates. No significant difference was observed among the wild type, DT7, DyukE and DT7E.

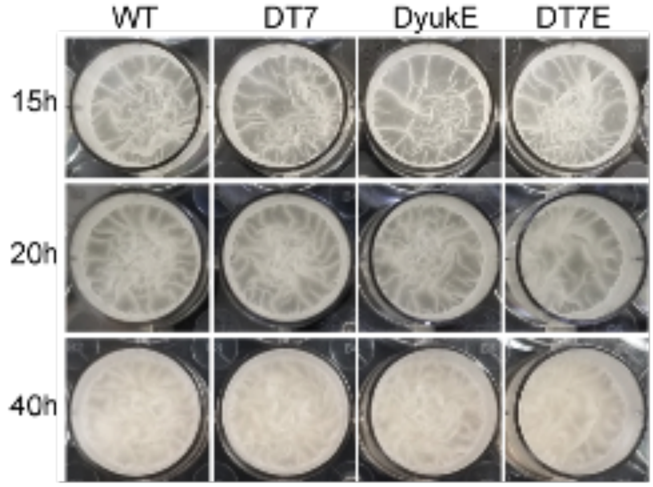


Fig. S3 Biofilm formation of *B. velezensis* SQR9 and the derived mutant strains. Biofilm formation by wild-type *B. velezensis* SQR9 and the mutants DT7SS, DyukE and DT7E was compared in MSgg medium, and no difference was observed during the whole period. The images shown here were taken at 15 h, 20 h and 40 h post inoculation.

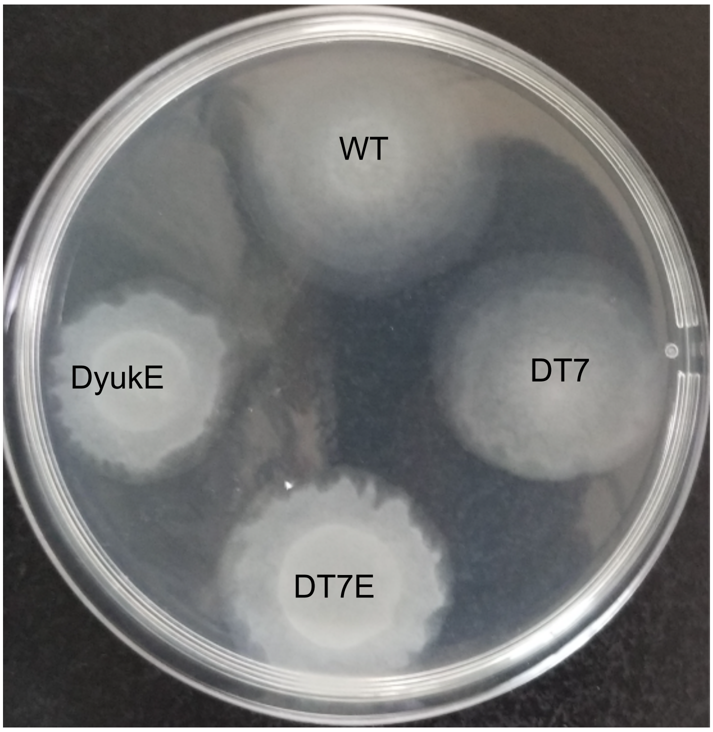


Fig. S4 Motility of *B. velezensis* SQR9 and the derived mutant strains. Cell motility of wild-type *B. velezensis* SQR9 and the mutants DT7, DyukE and DT7E was measured by a swarming assay. Bacterial cells were collected from LB medium cultures, washed twice with PBS and resuspended in swarming liquid medium. Equal volumes of cell suspension were inoculated on petri dishes containing semisolid LB medium. The petri dishes were incubated at 37°C for 6 h to allow the bacteria to swim. Then, the water content of the medium in the petri dishes was reduced with dry air to terminate the swimming process. Subsequently, the petri dish was incubated overnight at room temperature.

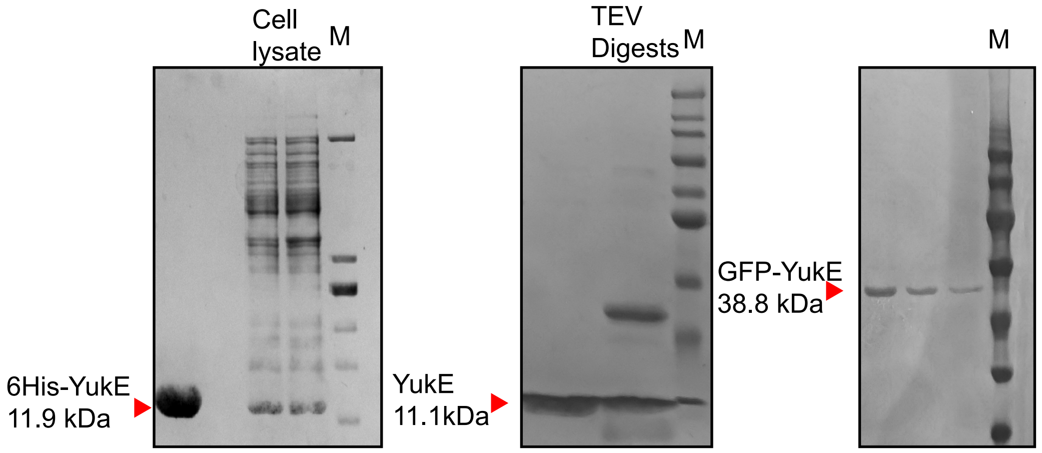


Fig. S5 Protein purification and SDS-PAGE of 6His-YukE, YukE and GFP-YukE. The coding sequences of the 6His-YukE and GFP-YukE fusion proteins were cloned into pET29a, and the recombinants were introduced into *E. coli* BL21(DE3) for expression. YukE without a His-tag was purified from 6His-tagged YukE linked to a TEV digestion site. The molecular weights of 6His-YukE, YukE and GFP-YukE were 11.9 kDa, 11.1 kDa and 38.8 kDa, respectively. The purity was higher than 90%.

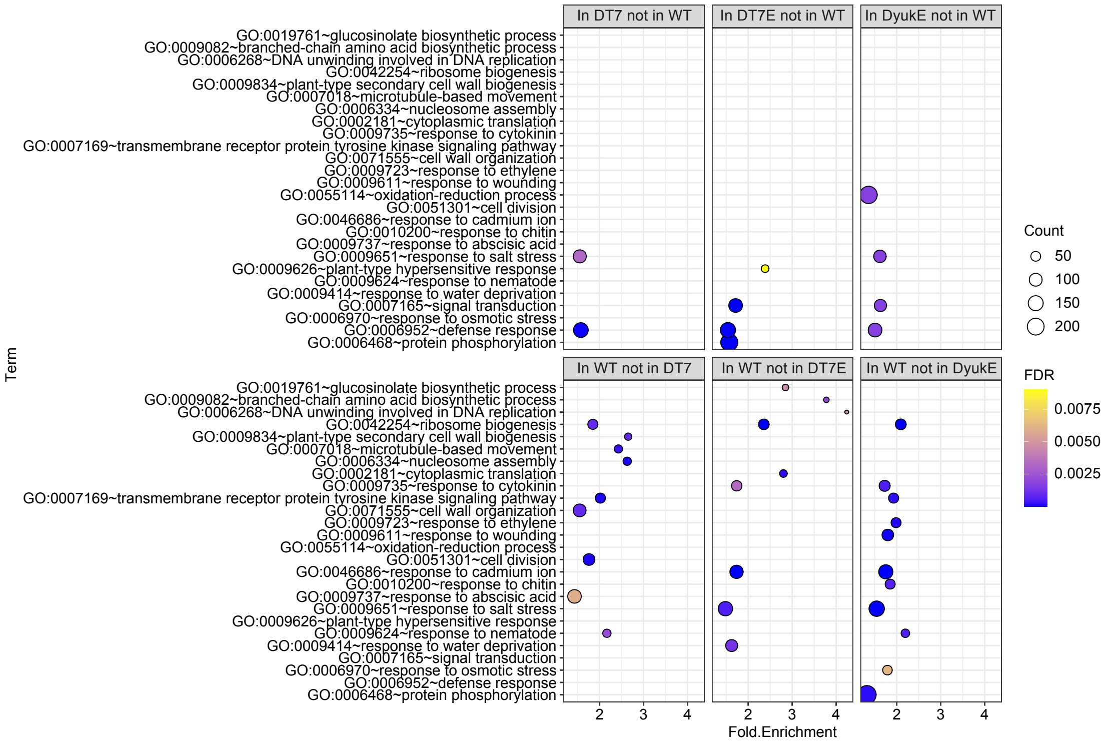


Fig. S6 Gene Ontology (GO) enrichment of Arabidopsis differentially expressed genes (DEGs) in the treatment with the *B. velezensis* mutants in comparison with that for the wild type. To identify the DEGs, the p value was calculated using a negative binomial distribution-based test, the FDR (p adjust) was calculated using BH (Benjamini-Hochberg) correction, and the DEGs were screened based on FDR < 0.05. DEGs for each mutant treatment were compared with those of the wild-type treatment, and the unique DEGs were subjected to GO analysis.

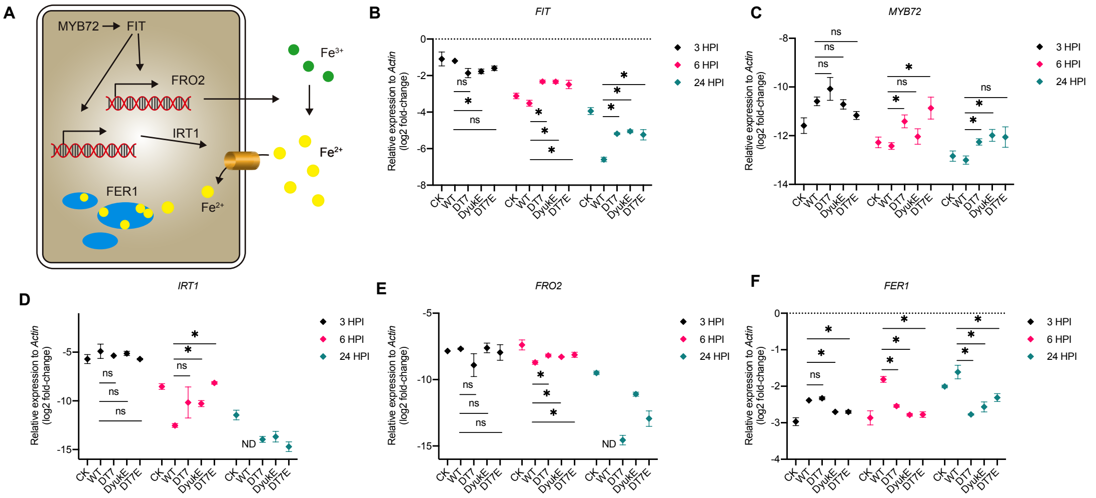


Fig. S7 Expression of plant iron acquisition genes measured by qRT-PCR. (A) The iron acquisition pathway in Arabidopsis. (B-F) Relative expression of *FIT* (B), *MYB72* (C), *IRT1* (D), *FRO2* (E) and *FER1* (F) in response to wild-type *B. velezensis* SQR9 and the mutants DT7, DyukE and DT7E. Expression in untreated plants was included as CK. Expression of *Actin* was measured as a reference. Each treatment included three independent replicates. Error bars indicate the standard error; “*” indicates a significant difference ($p<0.05$) based on Student’s t test, and “ns” indicates no significant difference. “ND” indicates not detected.

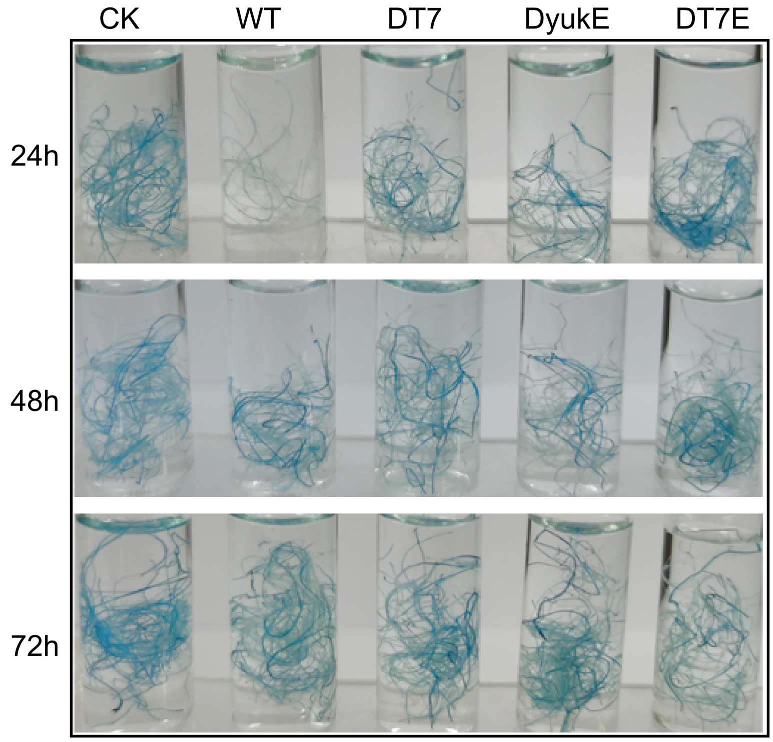


Fig. S8 Iron staining in plant roots. Arabidopsis roots cultured in 6-well plates were inoculated with wild-type *B. velezensis* SQR9 and the mutants DT7, DyukE and DT7E. After 24 h, 48 h and 72 h, the roots were collected, and the iron was stained with Perls' staining assay. Blue color in the photo indicates the iron in the roots. Each tube included roots from three plants, and each treatment included three independent replicates. The experiment was repeated twice and showed the same result.

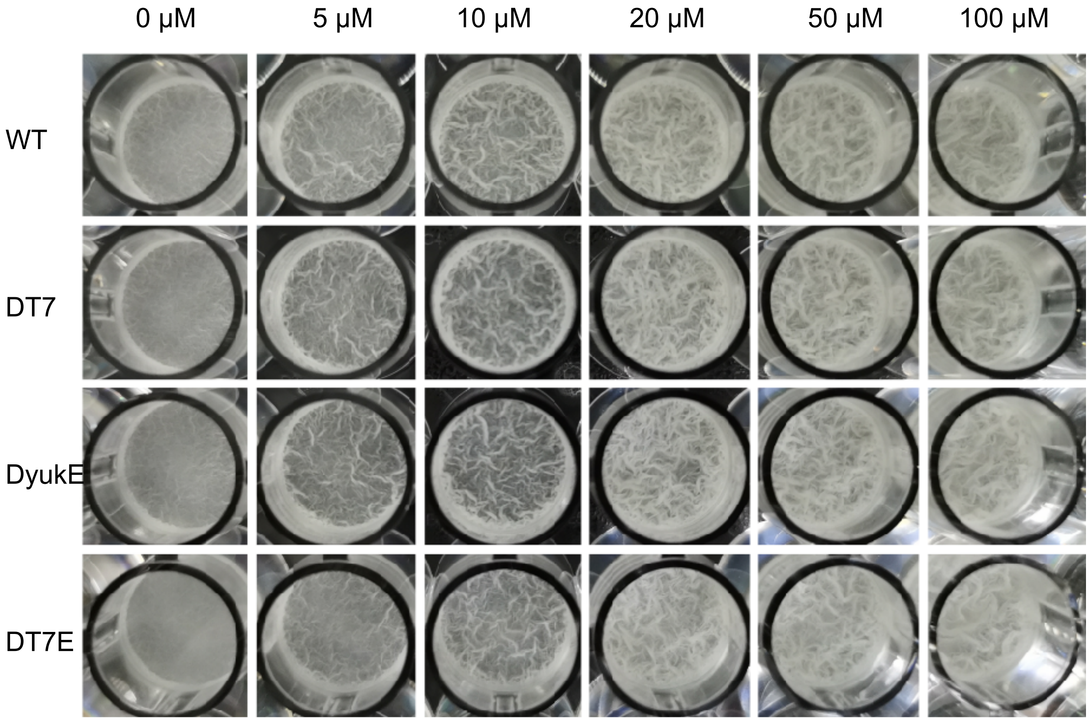


Fig. S9 Biofilm formation of *B. velezensis* SQR9 and the derived mutant strains with exogenous iron. A biofilm formation assay was performed with wild-type *B. velezensis* SQR9 and the DT7, DyukE, DT7E mutants in iron-free MSgg medium. Exogenous $FeSO_4$ was added at final concentrations of 0 μM, 5 μM, 10 μM, 20 μM, 50 μM and 100 μM. Three replicates were included for each treatment. The image shown was taken at 24 h post inoculation.

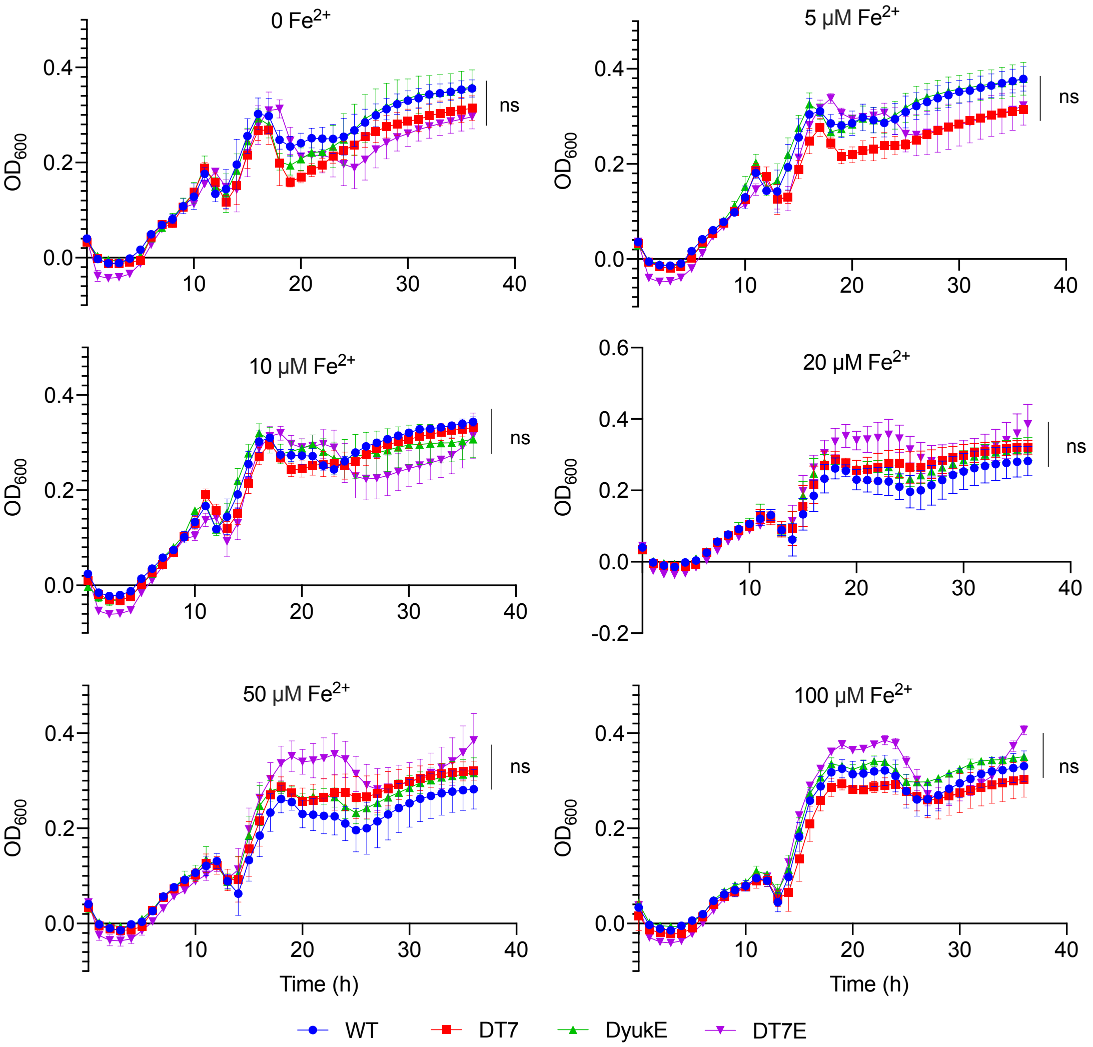


Fig. S10 Growth curve of *B. velezensis* SQR9 and the derived mutant strains in MSgg under different iron concentrations. Growth curves were measured by Bioscreen C. Wild-type strain *B. velezensis* SQR9 and the DT7, DyukE, DT7E mutants were grown in iron-free MSgg medium. Exogenous $FeSO_4$ was added at final concentrations of 0 μM, 5 μM, 10 μM, 20 μM, 50 μM and 100 μM. Six replicates were included for each treatment. Error bars indicate the standard error; “ns” indicates that no significant difference was found between treatments.

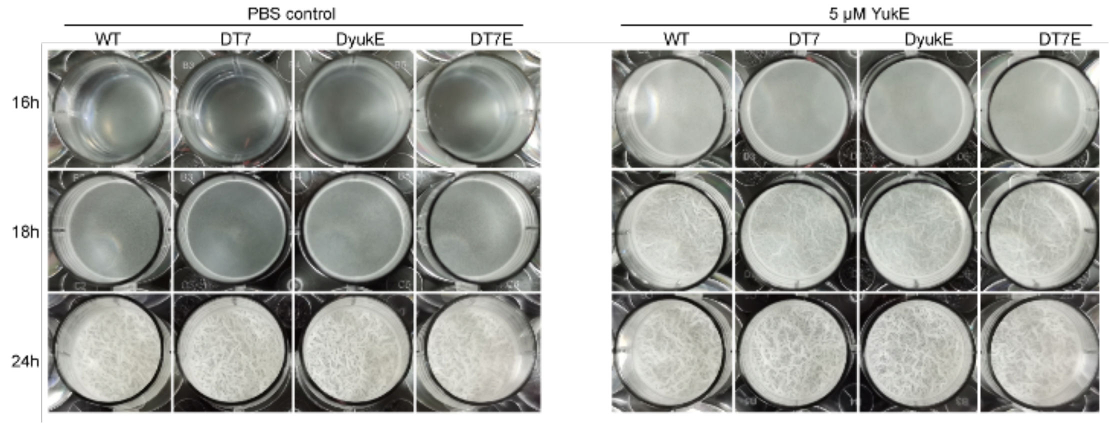


Fig. S11 Biofilm formation of *B. velezensis* SQR9 and the derived mutant strains with exogenous YukE. A biofilm formation assay was performed with wild-type *B. velezensis* SQR9 and the DT7, DyukE, DT7E mutants. Exogenous YukE was added at a final concentration of 5 μM. Three replicates were included for each treatment.

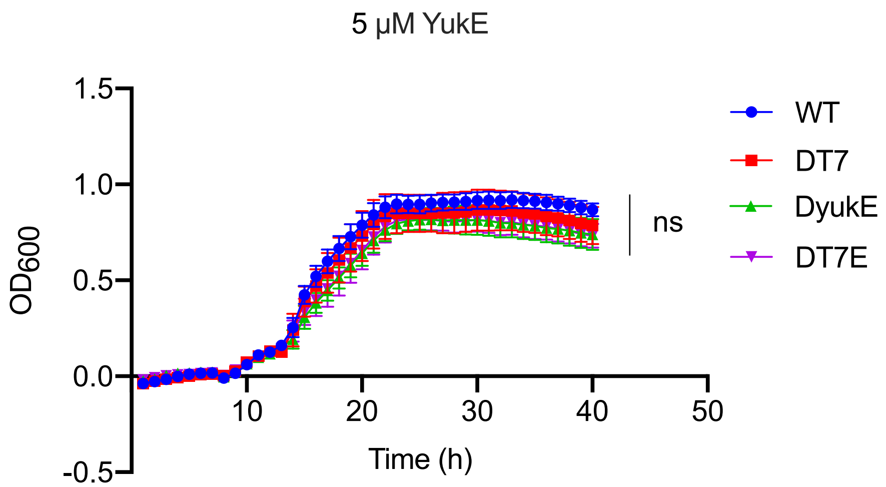


Fig. S12 Growth curve of *B. velezensis* SQR9 and the derived mutant strains with YukE. Growth curves were measured by Bioscreen C under the same culture conditions as the biofilm formation assay. Six replicates were included for each treatment. Error bars indicate the standard error; "ns" indicates that no significant difference was found between treatments.

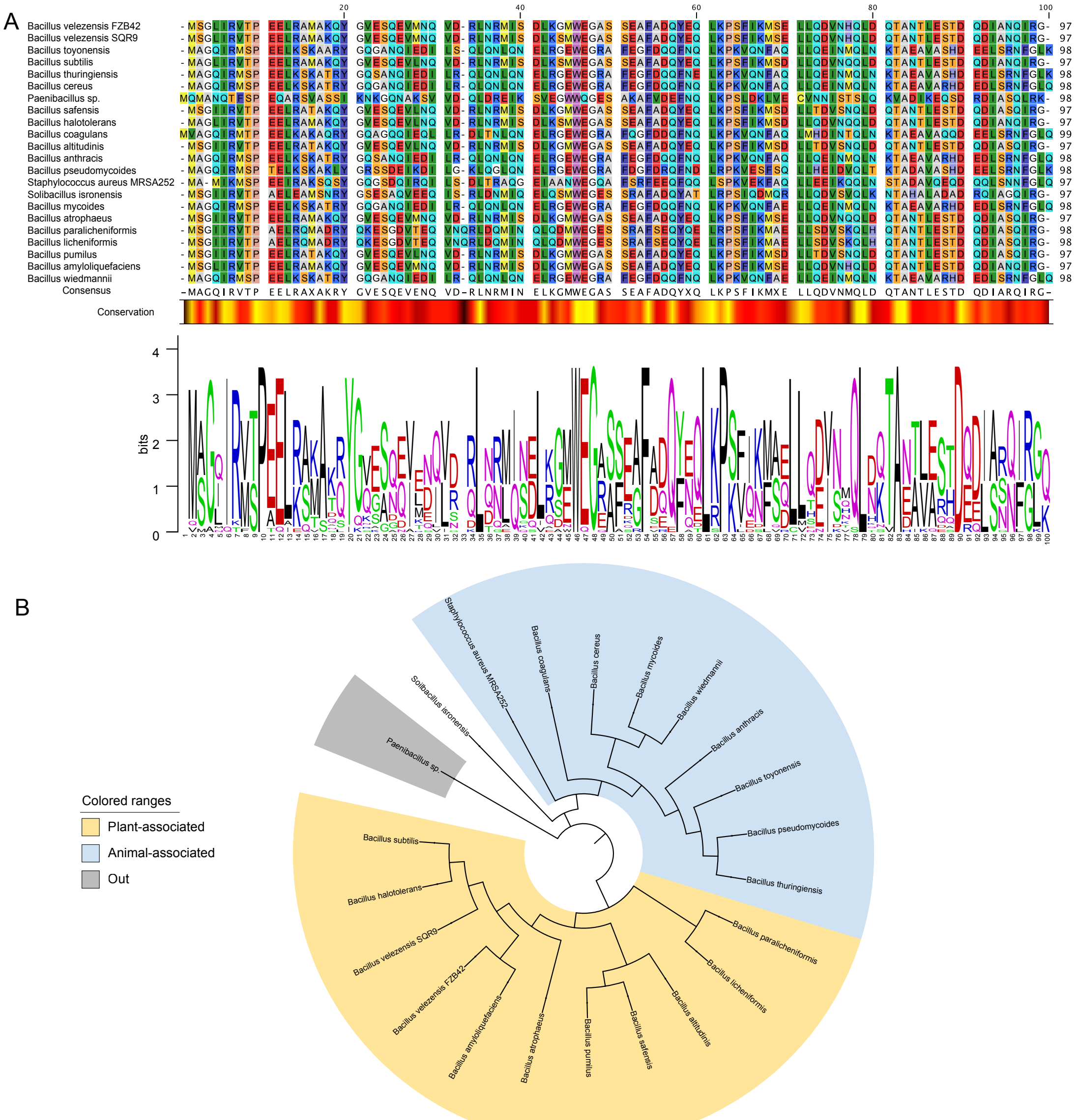


Fig. S13 Phylogenetic analysis of YukE in different *Bacillus* species. (A) Alignment of YukE. Twenty-two YukE sequences from *Bacillus* and closely related species were collected, and the natural niches of these bacteria were recorded. The alignment was performed with ClustalW algorithms using MEGAX. The conservation band shows the conservation at each amino acid position. Yellow indicates higher conservation, and red indicates lower conservation. (B) Phylogenetic tree for YukE. The phylogenetic analysis was inferred by using the maximum likelihood method based on the Tamura-Nei model. The initial tree for the heuristic search was obtained automatically by applying the neighbor-joining and BioNJ algorithms to a matrix of pairwise distances estimated using the maximum composite likelihood (MCL) approach and then selecting the topology with the superior log likelihood value. The analysis involved 22 nucleotide sequences. All positions containing gaps and missing data were eliminated. The phylogeny was tested by 500 bootstrap replications.

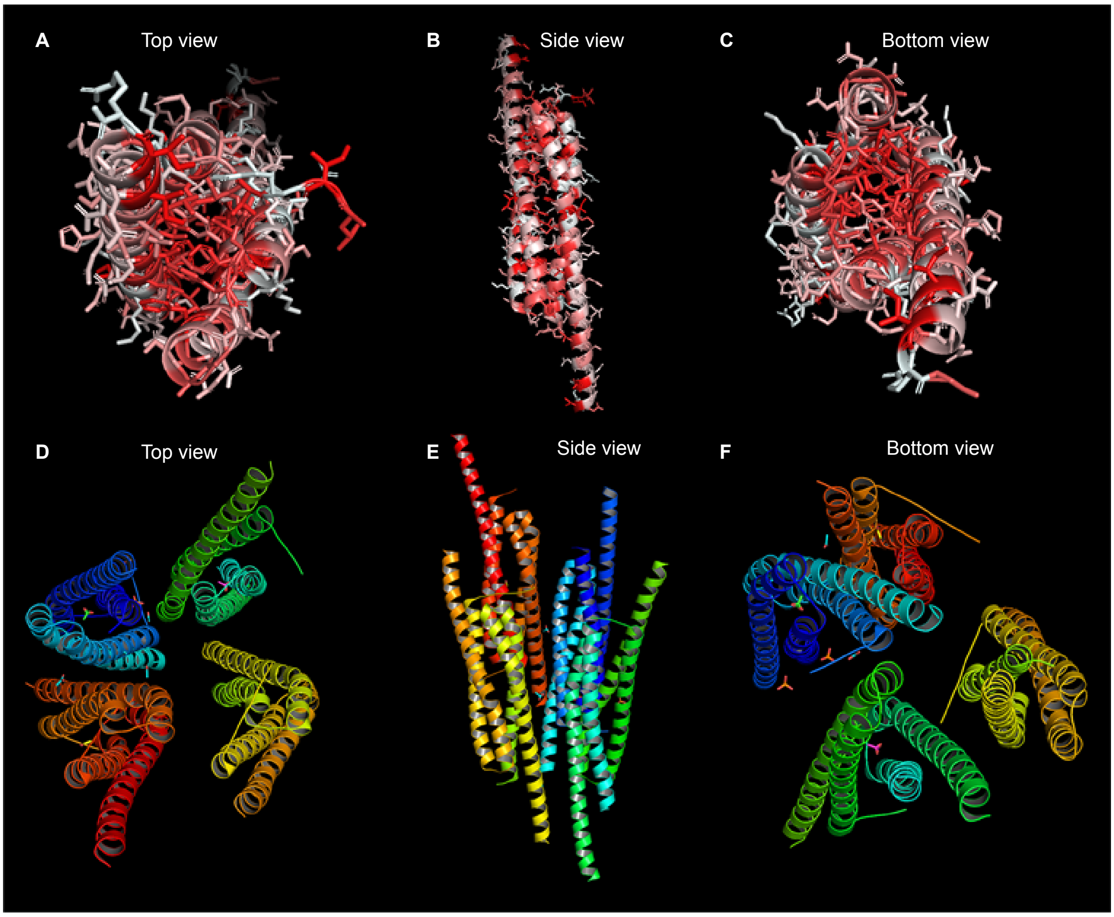


Fig. S14 Protein structure of YukE and EsxA. (A-C) Structure of YukE homodimer built by Swiss-model using EsxA of *M. tuberculosis* as template. The polarity of the residues was colored. Red indicated a higher polarity calculated by PyMOL (V. 2.4.1). (D-E) Asymmetric unit of EsxA from *Geobacillus thermodenitrificans.* Different color indicated different molecule.
